# Testing and overcoming the limitations of Modular Response Analysis

**DOI:** 10.1101/2024.08.21.608972

**Authors:** Jean-Pierre Borg, Jacques Colinge, Patrice Ravel

**Affiliations:** Université de Montpellier, Montpellier, France; Institut régional du Cancer Montpellier (ICM), Montpellier France; Institut de Recherche en Cancérologie de Montpellier (IRCM), Inserm U1194, Montpellier, France

**Keywords:** MRA, regression, network inference, Lack Of Fit, convex optimization

## Abstract

Modular Response Analysis (MRA) is an effective method to infer biological networks from perturbation data. However, it has several limitations, such as strong sensitivity to noise, need of performing independent perturbations that hit a single node at a time, and linear approximation of dependencies within the network. Previously, we addressed MRA sensitivity to noise by revisiting MRA as a multilinear regression problem. Here, we provide new contributions to complement this theory. First, we overcame the need of perturbations to be independent, thereby augmenting MRA applicability. Second, using analysis of variance (ANOVA) and lack of fit tests, we assessed MRA compatibility with the data and identified the primary source of errors. If nonlinearity prevails, we propose a polynomial extension to the model. Third, we demonstrated how to effectively use the prior knowledge of the network studied. Finally, we added these innovations to our R software package MRARegress to provide a complete, extended theory around MRA and to facilitate its access by the community.

## Introduction

The study of biological systems is inherently complex because they are governed by intricate molecular interactions that orchestrate the cell and tissue vital functions and responses to external stimuli. Therefore, the inference of biological networks is critical to understand homeostasis and diseases. Depending on the study granularity level, biological networks may involve genes, proteins, metabolites, or larger structures, such as protein complexes or even a subnetwork that would be regarded as a single object. Combinations are possible with some network participants, such as individual molecules, whereas other participants could be larger structures that do not need to be modeled in details. For concision, we name these unresolved larger structures, ‘modules ‘.

In addition, to determine the activity of each network participant, a quantitative measure must be obtained in the various tested experimental conditions. In the case of a gene, transcriptional abundance is a natural measure, but it could also be the methylation level of its promoter. In the case of a protein, its direct abundance or activation status (*e*.*g*., phosphorylation level) is an obvious choice. In the case of a module, the chosen measure must represent its overall activity.

Several methods can be used to infer biological networks from the experimental measurements of their component activity (Mekedem *et al*., 2022). These methods can be broadly categorized as follows:

- Data-based methods: such as statistical correlations (e.g. Pearson, Spearman, or Kendall (Alberts, 2002)), Mutual Independence (e.g., ARACNE (Margolin *et al*., 2006), CLR (Faith *et al*., 2007) or MRNET (Meyer *et al*., 2007)), probabilistic graphical models (e.g., GeneNet (Schaffter *et al*., 2011)), Bayesian (Friedman and Koller, 2003), (Hill *et al*., 2012) and regression methods (e.g., TIGRESS (Haury *et al*., 2012)).
- Machine learning methods (Random Forest (Huynh-Thu *et al*., 2010), Support Vector Machine (Brouard, 2013), neural networks (Cardot *et al*., 2013)). These methods are efficient, but require extensive datasets for training and validation.
- Perturbation-based methods, including the classical Modular Response Analysis (MRA) (Kholodenko *et al*., 2002)), built using Ordinary Differential Equations (ODEs) that describe the network dynamics. MRA is an effective method to infer networks, including when granularity may vary (*i*.*e*., the network includes modules). To use MRA, the network dynamics must have reached a steady-state when the experimental measures are acquired.

This article focuses on the extension of the MRA approach to embed it in linear and polynomial regressions, and to exploit the available prior knowledge on some interactions in the network to be inferred. The first MRA developments were proposed by Boris Kholodenko in his seminal paper (Kholodenko *et al*., 2002). Subsequent enhancements incorporated noisy data integration and statistical estimation of connectivity coefficients, using the maximum likelihood principle (MLMSMRA (Klinger and Blüthgen, 2018), (Bosdriesz *et al*., 2018)) or by estimating the confidence interval of the parameters by Monte Carlo analysis (Andrec *et al*., 2005), (Santos *et al*., 2007).

Other authors adopted a Bayesian approach (Santra *et al*., 2013), (Santra *et al*., 2018), leveraging the distribution of prior knowledge on connectivity coefficients and partial network topology, including their sparsity, extracted from the literature or reference databases, such as STRING (Szklarczyk *et al*., 2011). In our recent work, we extended MRA by expressing it as a multilinear regression problem (Borg et al., 2023) where the connectivity coefficients, derived from the regression residual variance, and the model parameter were estimated in parallel. This methodology, which we named MRARegress, allowed us to define a whole family of MRA methods, one method for each regression algorithm essentially (e.g., LASSO, STEP, least squares, Random Forest).

One of the important issues, raised by most of the previously quoted authors, is MRA sensitivity to measurement noise and perturbation levels, commonly known as signal-to-noise ratio (Andrec et al., 2005), (Thomaseth et al., 2018), (Borg et al., 2023). This highlights the need to consider nonlinearity.

Comparative analysis of the classical MRA method (Kholodenko *et al*., 2002), some methods based on Mutual Independence (Margolin *et al*., 2006) and MRARegress demonstrated MRARegress superior performance in the presence of noisy data and medium to large networks (10 to 1000 nodes) (Borg *et al*., 2023).

Here, we enriched MRARegress functionalities by proposing holistic solutions to its constraints that we then validated across networks with known dynamics and extensive simulations. Specifically, following a review of the principles and prerequisites for applying MRA:

- We optimized the use of the available measurement data, enabling the consideration of technical and biological replicates to estimate the parameter confidence intervals.
- We introduced a Lack Of Fit (LOF) test to assess nonlinearity versus measurement noise and determine the linear model relevance.
- For cases when the model is not relevant, we detailed an extension of MRARegress that accommodates nonlinearity by incorporating second-order terms, thereby harnessing potential synergistic effects between variables.
- For cases when prior knowledge of the studied network is available, we showed that MRARegress, coupled with convex optimization, effectively incorporates this knowledge.

These methodological developments are included in a R software package, that we also named MRARegress, and that provides user-friendly and free access to our new MRA algorithms for the scientific community.

## Methods

In MRA, every node within a network is sequentially perturbed and the network response to these perturbations is analyzed. Each node must be associated with a measurable quantity related to its activity: abundance or state (e.g. phosphorylation level of a protein or expression level of a gene). Perturbations may include administering a drug, siRNA or shRNA that completely inhibits, reduces or increases the expression of a gene. These perturbations are referred to as Knock Out (KO), Knock Down (KD) or Knock Up (KU), respectively.

### 1 Mathematical aspects

#### 1.1 Expression of Δ*X*_*i*_ /based on Δ*X*_*k*_, *k*≠*i*

All the measures associated with the *N* modules at a given time will be represented by a vector *X* with *N* dimensions. Thus, *X*_*i*_(*t*) corresponds to the measurement associated with node *i* at time *t*. Because of the interconnection of the modules, *X*_*i*_ depends on the other *X*_*j*,_ (*j*≠*i*).

All *X*_*i*_, *i* ∈ ⟦1, *N*⟧also depend on a set of *M* parameters *p*_*k*_, *k* ∈ ⟦1, *M*⟧(identified by a vector *P*) and the time *t*.

In a very general way, this vector X is the solution of the system of differential equations:

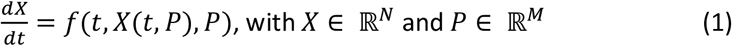

The system of differential equations may not be a first-order system, but all systems can be rewritten as first-order differential equations by introducing additional variables.

A fundamental assumption of MRA is that there is a set of parameters, referred to as *P*_0_, a neighborhood of *P*_*0*_ (noted *V(P*_*0*_)) and a time *t*_*0*,_ beyond which the system reaches a stable and stationary state, which will be noted *X(P*_*0*_). In other words, beyond *t*_*0*_, *f* no longer depends on the time t and the *dX*/*dt* derivative is null. For networks that support this hypothesis, (1) becomes:

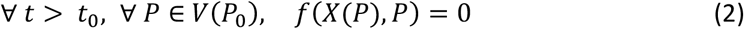

Therefore, MRA cannot be used when the network dynamics are periodic or pseudo-periodic.

MRA estimates the gradient of the function *f* and the gradient components represent the interactions of the module network. They are called connectivity coefficients and are the unknowns of the problem.

Function *f*, of ℝ^*N*+*M*^ in ℝ, is usually not known. Its gradient can be approached with the following development.

Assuming the function *f* of class *C*^1^, it can be demonstrated, using the implicit function theorem (Couty and Ezra, 1970) and Taylor ‘s first order development, that for each component *X*_*i*_ of vector *X*, there is a function *φ*_*i*_ which depends on other *X*_*j*_ components and parameters, such as:

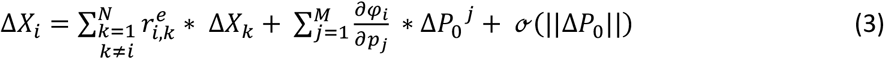

Functions *φ*_*i*_ have continuous partial derivatives in the neighborhood of (*X*(*P*_0_)), *P*_0_). In this equation:

- Δ*X*_*j*_ = *X*_*j*_(*P*_0_ + Δ*P*_0_) − *X*_*j*_(*P*_0_), ∀*j* ∈ ⟦1, *N*⟧, represents the modification of the *j* module expression measurement when applying a Δ*P*_0_ perturbation from the stationary state *P*_*0*_.
- 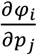 is calculated at the point *P*_0_. The theory tells the existence of these functions *φ*_*i*_, but not how to compute them. The following paragraphs will show how to get rid of these terms.
- 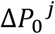 represents the j^th^ coordinate of Δ*P*_0_ in ℝ^*M*^.
- ℴ(|| Δ*P*_0_||) is a function related to Taylor’s remainder. It measures the nonlinearity of *f*. It is 0 if the Δ*P*_0_ perturbation is null or if *f* is a linear function. Conditions will be examined to make this amount as small as possible.
- 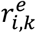, called exact connectivity coefficient, represents the action of node *k* on node *i*.

If 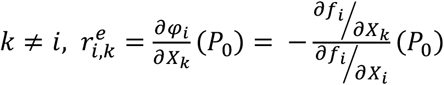. By definition, 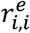 has no meaning. It is given the value -1 to simplify the subsequent calculations.

#### 1.2 Assumption of the independence of perturbations

Now, let us assume that:

*M* = *N, p*_*i*_ influences ONLY *X*_*i*_ (in other words: 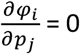 if *j*≠*i* and 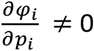), AND perturbations Δ*P*_0_ can be achieved that affect only the parameter *p*_*i*_, ∀*i* ∈ ⟦1, *N*⟧.

This strong hypothesis, both from a mathematical and biological point of view, was made implicitly or explicitly in various articles on MRA (Kholodenko *et al*., 2002), (Jimenez-Dominguez *et al*., 2021), (Mekedem *et al*., 2022). Throughout this article, this hypothesis will be designated as the ‘Assumption of Independence Of Perturbations ’ (AIOP). In these conditions, equation (3) becomes:

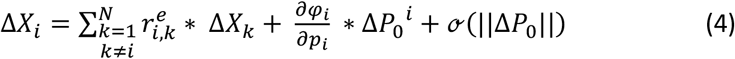

See Figure 1 for a geometric interpretation of this equation.

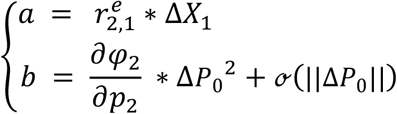

**Figure 1:**
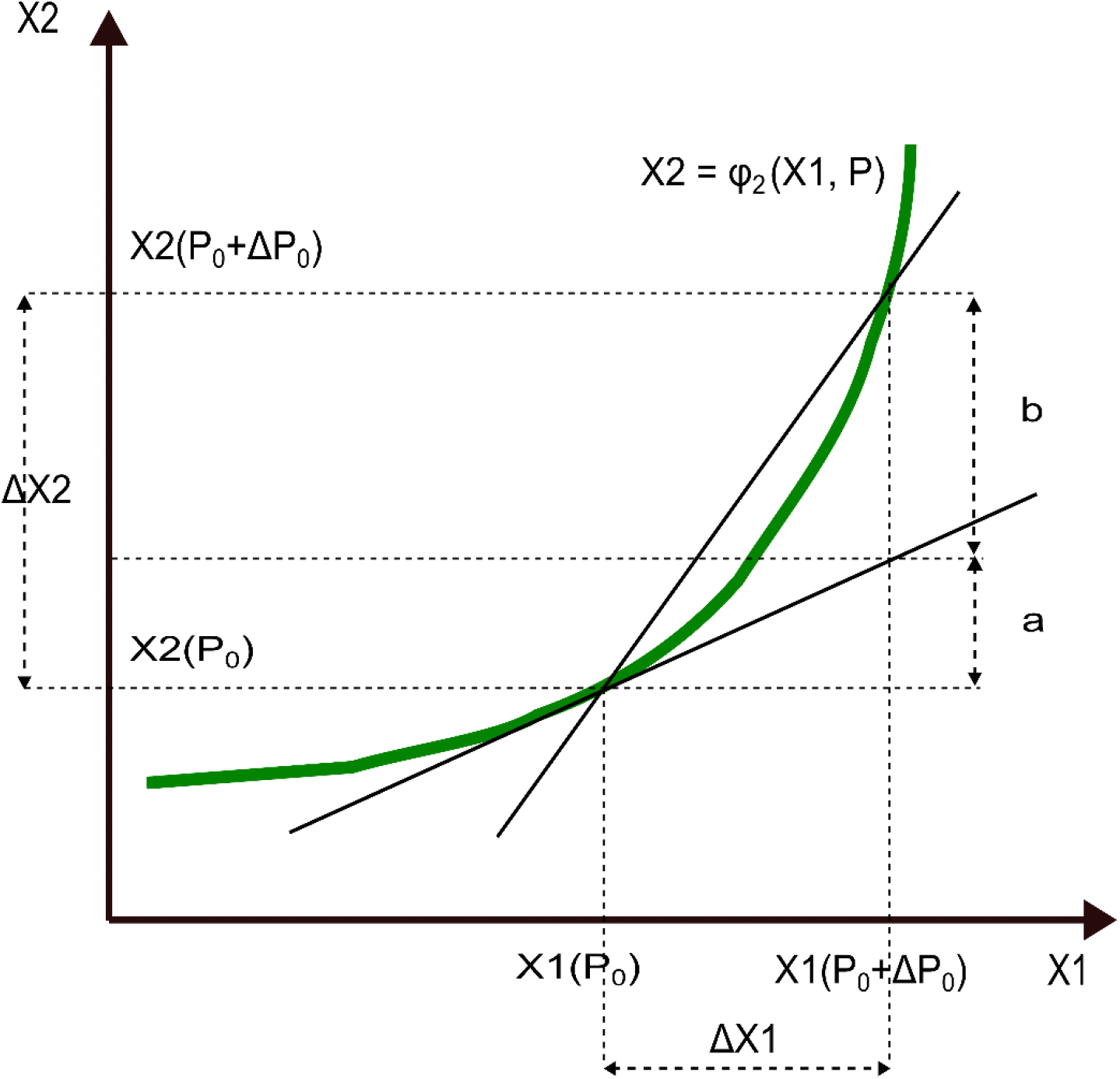
Geometric interpretation of equation (4) for a 2-node network

By neglecting the remainder ℴ(|| Δ*P*_0_||), which is acceptable for small perturbations, and by remembering that 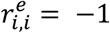, the system of approximate equations is obtained

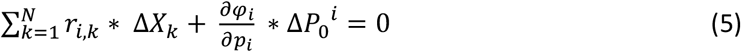

(note that 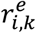 have become *r*_*i,k*_, connectivity coefficients).

If one chooses a Δ*P*_0_ perturbation that acts ONLY on the parameter *p*_*j*_, (*j*≠*i*), therefore on the node *j*, which is possible according to the AIOP (perturbation designated by *q*_*j*_), the coordinates of *q*_*j*_ on the other parameters *p*_*k*_, with *k*≠*j* are null.

This leads to a system of N - 1 equations to N – 1 unknowns *r*_*i,k*_, *k*≠*i*, if N -1 perturbations of this type can be performed.

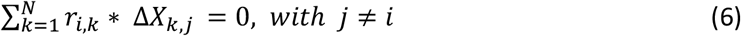

In this system of equations, Δ*X*_*k,j*_ = *X*_*k*_(*P*_0_ + *q*_*j*_) − *X*_*k*_(*P*_0_) and *r*_*i,i*_ = −1. The solution of this system allows computing the coefficients *r*_*i,k*_ if the rank of the matrix (Δ*X*_*k,j*_), *k* and *j*≠*i* is N-1.

By giving *i* the successive values 1 to N, all coefficients *r*_*i,j*_ of the connectivity matrix, called “r ”, can be obtained. **The coefficient *r***_***i***,***j***_ **is an approximate value of** 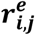 and will be called connectivity coefficient. It also represents the action of node j on node i.

These remarks are illustrated in Figure 2.

**Figure 2:**
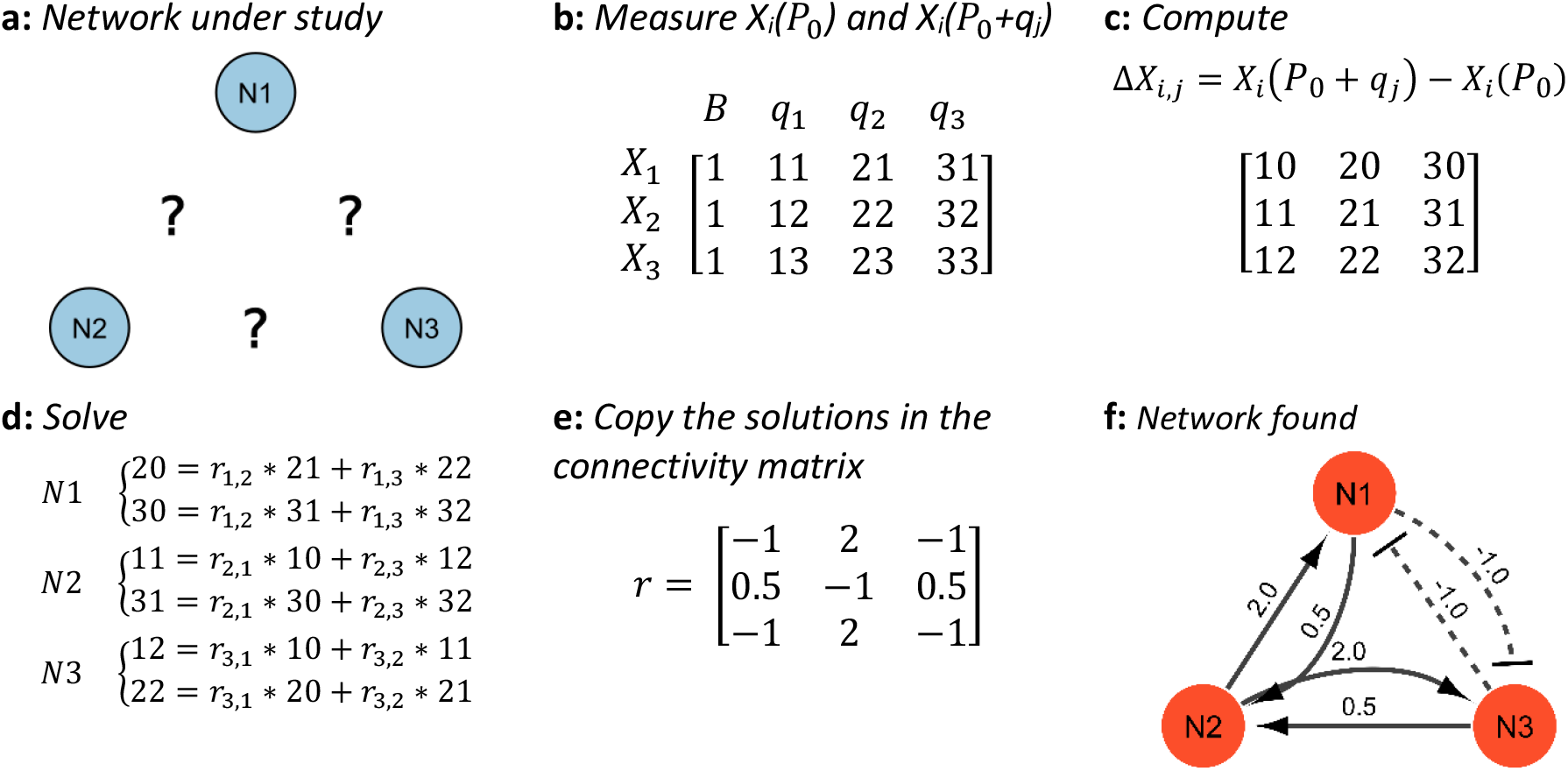
Illustration of the MRARegress method (a). In this 3-node network, we want to know the interactions between nodes. (b). Numerical example of the actions to be performed (displayed matrix): -measure the expression of the three modules in the non-perturbed (called basal, B) state: X_1_(P_0_), X_2_(P_0_), X_3_(P_0_)) (column 1), -apply perturbation q_1_ to module 1. Wait until the steady state is reached, measure the same expressions again: X_1_(P_0_+q_1_), X_2_(P_0_+q_1_), X_3_(P_0_+q_1_)) (column 2), -repeat by applying perturbation q_2_ to module 2 (column 3) and perturbation q_3_ to module 3 (column 4). (c). Then, the values ΔX_1,1_ = X_1_(P_0_ + q_1_) − X_1_(P_0_), ΔX_2,1_ = X_2_(P_0_ + q_1_) − X_2_(P_0_), ΔX_3,1_ = X_3_(P_0_ + q_1_) − X_3_(P_0_) are computed to find the action of N2 and N3 on N1. The other values ΔX_i,j_ are calculated to find the actions on the other nodes. (d) To determine the action of nodes 2 and 3 on node 1, solve (the unknowns are r_1,2_ and r_1,3_)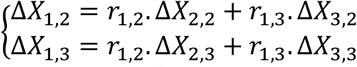 Similarly solve 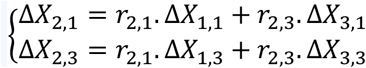 to determine the action of nodes 1 and 3 on node 2 and 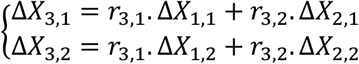.to determine the action of nodes 1 and 2 on node 3. (e) The Connectivity Matrix “r ” is obtained. We remember that r_i,i_ = −1, for i ∈ {1,2,3}. (f). These interactions can be represented as: 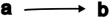 a amplifies the action of b (ri,j>0) 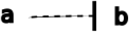 a inhibits the action of b (ri,j<0).

If the AIOP is **not true**, equation (3) cannot be simplified. By neglecting the remainder ℴ(|| Δ*P*_0_||), equation 7 is obtained:

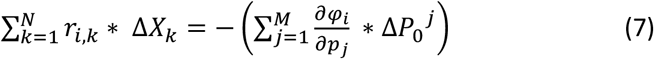

This equation can be applied to *Q* perturbations *q*_*m*_, *m ϵ* ⟦1, *Q*⟧. For each value of *i ϵ* ⟦1, *N*⟧, equation (7) becomes a system of *Q* linear equations:

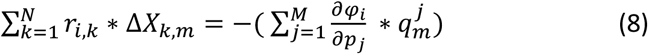

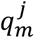 is the j^th^ coordinate of perturbation *q*_*m*_ in ℝ^*M*^ and Δ*X*_*k,m*_ = *X*_*k*_(*P*_0_ + *q*_*m*_) − *X*_*k*_(*P*_0_).

This system of *Q* equations (8) has *N*− 1 + *Q* ∗ *M* unknowns in general:

- 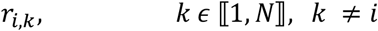
- 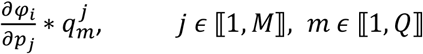

Without additional hypothesis on perturbations, this system cannot be solved because there are more unknowns than equations. Conversely, if the AIOP is *partially* verified (it is known that some 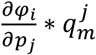 are null for all *j*), this will reduce the number of system (8) unknowns. Precisely, for each *i* call *IND*_*i*_ the set of perturbations *m* such as 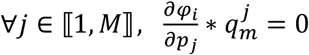. If the rank of the matrix (Δ*X*_*k,m*_), *k* ∈ ⟦1, *N*⟧and *m* ∈ *IND*_*i*_ is *N-1* for all *i*, the system (8) will have a solution.

#### 1.3 Principle of the LOF test

For a node *i* of the network, equation (6), in which the remainder ℴ(|| Δ*P*_0_||) is not neglected, can be written as follows, because *r*_*i,i*_ = −1:

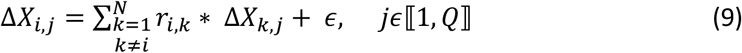

Errors due to nonlinearity ℴ(|| Δ*P*_0_||) and measurement noise have been grouped in the term *ϵ*.

In the presence of replicates, under certain regularity assumptions (Saporta, 1990), the LOF test can be applied to a multilinear regression (Mason *et al*., 2003). This technique allows us, using replicates, to estimate for each node the pure error variance (due to measurement noise only), as shown in Figure 3.

**Figure 3:**
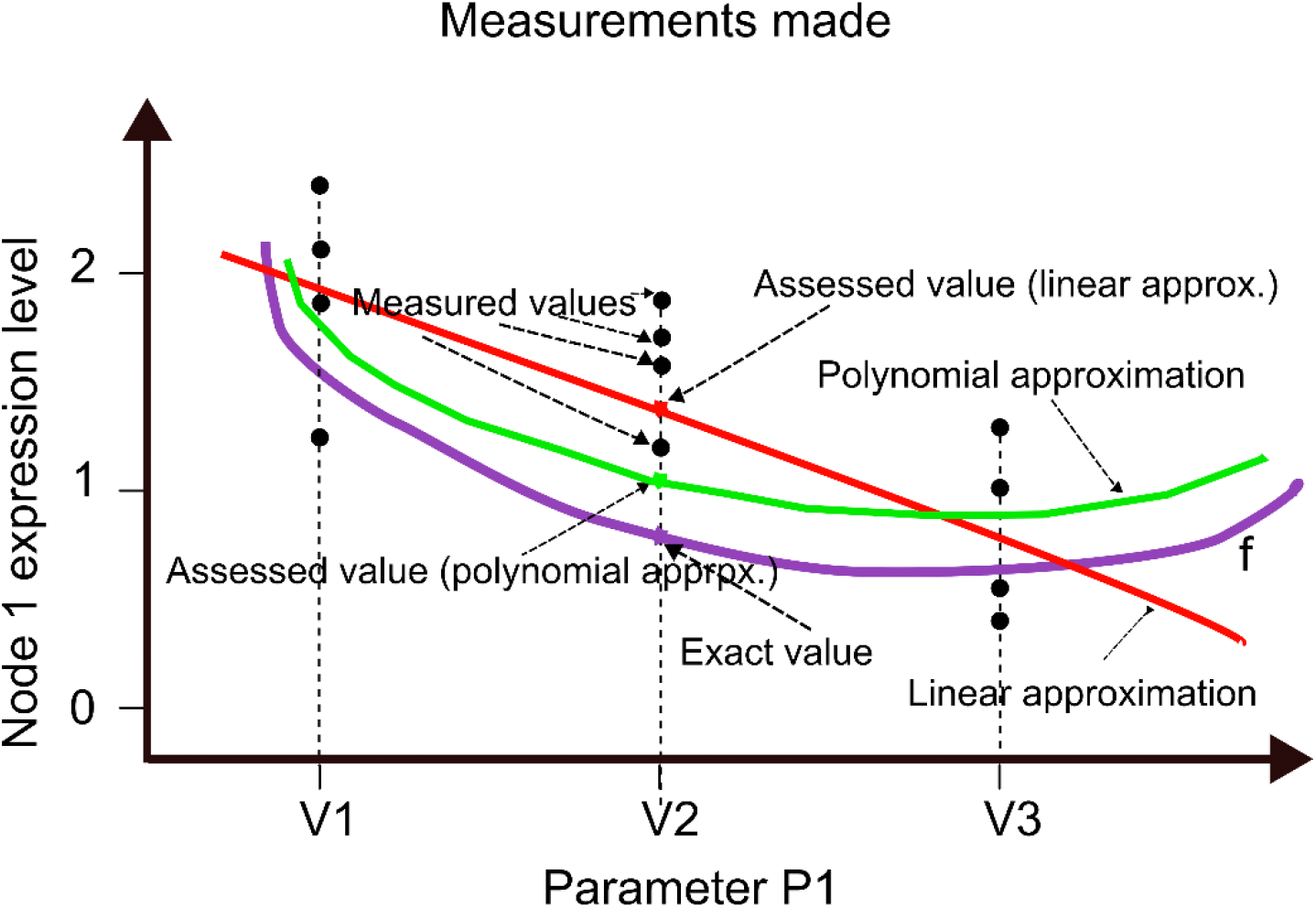
Illustration of the estimation of connectivity coefficients using linear or polynomial regression.

For each node *i*, Q measurements were performed (perturbations including replicates, Q = 12 for P_1_ in Figure 3). To simplify, it was assumed that the numbers of replicates and perturbations were the same for all nodes, but this is not mandatory and the MRARegress package considers the exact values.

##### First ANOVA, regarding Regression

Following the Mason’s notations, *TSS* is the Total Sum of Squares, *SSR* is the Sum of Squares due to Regression, *SSE* is the Sum of Squares Error, and 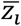 is the mean of the results obtained for this node. 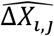 correspond to the values of equation (6) in § 1.2, in which the unknowns *r*_*i,j*_ have been replaced by their values estimated by linear regression.

The MRARegress approach is associated with a zero-intercept regression model (see eq. (6)). In this case, the TSS associated with the model breaks down as follows:

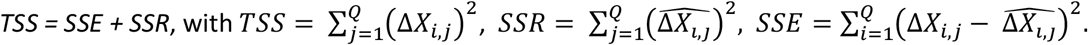

The corresponding degrees of freedom (*df*) are Q, N-1, and Q-N+1 respectively. ANOVA can be performed for this regression parameter test (Saporta, 1990).

##### Second ANOVA, regarding LOF

By using replicates, it is possible to determine the SSE pure error (*SSE*_*P*_: noise error of the measurements) and the error caused by the model mismatch (*SSE*_*LOF*_: component due to LOF test error).

The number of different perturbations is noted as *m* (m = 3 for P_1_ in Figure 3) and for each perturbation t, *n*_*t*_ is the number of the corresponding replicates (4 for all of them in Figure 3). Of course,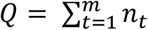. Let’s call Δ*X*_*i,t,k*_ the k^th^ measure corresponding to the perturbation *t* and 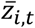 the mean of the measures corresponding to that perturbation.

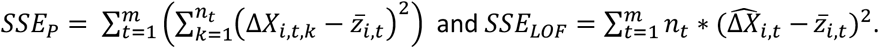

The LOF test is a Fisher’s test of the estimated variances 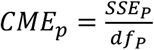 and 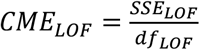 where the degrees of freedom are 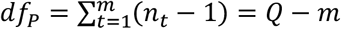 and *df*_*LOF*_ = (*Q* − *N*+ 1) − (*Q* − *m*) = *m* − *N*+ 1, respectively.

If the *CME*_*LOF*_/*CME*_*p*_ ratio is large (p-value < 0.05), this test rejects the null hypothesis “the residual error can be explained by the measurement noise ” and concludes that there is a significant effect of the model nonlinearity, relative to the measurements noise. Otherwise, the linear model is considered acceptable if the noise level, estimated by the replicates, is also acceptable.

#### 1.4 Extension to polynomial regression (order 2)

If the linear model is not acceptable, the reasoning that led to equation (3) could be extended by keeping the previous assumptions, but by being more demanding concerning the regularity of function *f*:

*f is now a class C*^2^ *function of* ℝ^*N*+*M*^ *in* ℝ, *functions φ*_*i*_ *have continuous first- and second-order partial derivatives in the vicinity of* (*X*(*P*_0_)), *P*_0_).

Using again the implicit functions theorem (Couty and Ezra, 1970) and Taylor’s development to order 2 this time, it can be demonstrated that for each *X*_*i*_ component of vector *X* in the vicinity of *P*_0_:

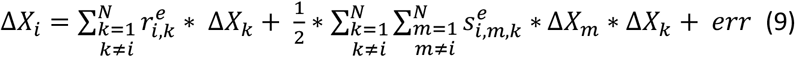

In this equation, as with equation (3):

- Δ*X*_*j*_ = *X*_*j*_(*P*_0_ + Δ*P*_0_) − *X*_*j*_(*P*_0_), ∀*j* ∈ ⟦1, *N*⟧, represents the modification of the *j* module expression measurement, as written in equation (3).
- 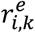, is still the exact connectivity coefficient (which reflects the action of node *k* on node *i*).
- 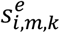 represents the influence of quadratic terms 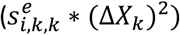 and quadratic cross terms 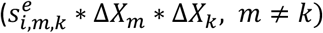
- The error *err* is determined by the amplitude of perturbations || Δ*P*_0_||.

Taylor’s formula tells that *err* is a negligible function compared with || Δ*P*_0_||, which is written, with the Landau notation (little-o), *err* = ℴ(‖ Δ*P*_0_‖). It corresponds to a first order error.

It can also be shown that if the function *f*, which describes the dynamics of the system and was defined in equation (2), is of the form *f*(*X*(*P*)) = 0 and not *f*(*X*(*P*), *P*) = 0, then the error is even smaller. This is a second-order error: *err* = ℴ(‖ Δ*P*_0_‖^2^).

For each *i*, equation (9) reveals N-1 numbers 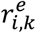 and 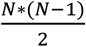 numbers 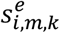 because 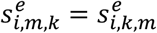. It is a total of 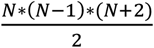 unknowns.

As in the linear case, by applying Q perturbations *q*_*p*_, with *p* ∈ ⟦1, *Q*⟧and neglecting an error of order 1 or 2, the approximate coefficients *r*_*i,k*_ and *s*_*i,m,k*_ can be found by solving the system of linear equations:

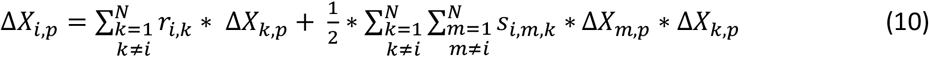

with *p* ∈ ⟦1, *Q*⟧and Δ*X*_*j,p*_ = *X*_*j*_(*P*_0_ + *q*_*p*_) − *X*_*j*_(*P*_0_), ∀*j* ∈ ⟦1, *N*⟧, if the system rank is sufficient.

### 2 Methods used to evaluate results

The purpose of the methods to measure module network inference is to calculate, as accurately as possible, the real connectivity coefficients 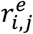, because they correspond to the interactions between modules in the network. To evaluate the new developments described in this article, the discovered connectivity matrix “ *r* ” and the real matrix “ *r*^*e*^ ” were compared by means of scores that depend on what is known about “*r*^*e*^ ” (see § 0).

#### 2.1 Test networks where 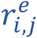 exact values are known

This is the case when the system dynamics are known (partly from the work by B.N. Kholodenko) or can be simulated (FRANK Network Generator, named ‘FRANK ‘). For these networks,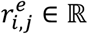. The chosen score is the distance (according to the L2-norm) between the exact connectivity matrix 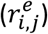 and the computed connectivity matrix (*r*_*i,j*_): *d* (*r*^*e*^, *r*).

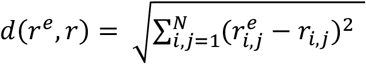. Smaller distances indicate better results.

#### 2.2 Test networks for which only the existence or absence of interactions between nodes is known

Here,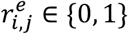. This is the case of Boolean networks, e.g. DREAM Challenge 4 (DC4). The Euclidean distance does not make sense in this case. Therefore, after digitizing the matrix “ *r* ” (by comparing |*r*_*i,j*_| to a threshold (Th) and obtaining an *rdig* matrix), a confusion matrix is computed from matrices (*rdig*) and (*r*^*e*^), from which two indicators are obtained: Sensitivity 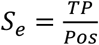 and Specificity 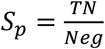, where:

– TP (True Positive): number of *r*_*i,j*_ estimated at 1 and actually worth 1.
– TN (True Negative): number of *r*_*i,j*_ estimated at 0 and actually worth 0.
– Pos: number of 1 in the solution.
– Neg: number of 0 in the solution. Pos + Neg = N.(N-1), where N is the number of nodes, because diagonal elements are known and are not evaluated (they are -1).

Comparing results from two values is not convenient. Furthermore, the choice of threshold for digitization (Th) exerts opposing influence on these ratios: decreasing Th increases the sensitivity and decreases specificity, and vice versa. To circumvent this issue, it is customary to construct a Receiver Operating Characteristic (ROC) curve that represents *Se* against 1 − *Sp* by varying Th (Fig. 4).

**Figure 4:**
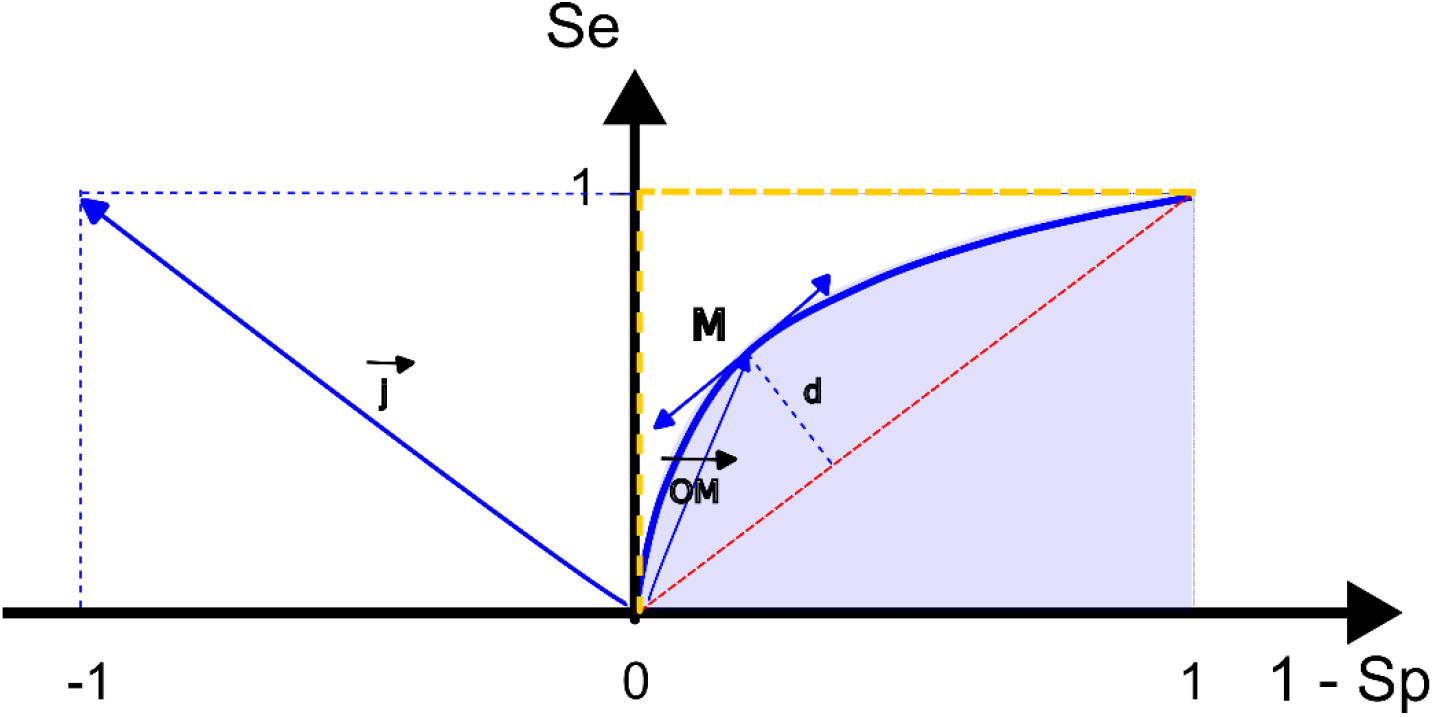
ROC curve and Distance to Diagonal 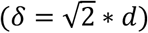. δ is equivalent to AuROC, but less time consuming.Reminder: all ROC curves intersect at coordinate points (0.0) and (1.1). In the instance of a purely random method (i.e., it randomly decides whether node j influences node I or not), this curve aligns with the bisector in red on the figure. The perfect detection corresponds to the dashed yellow curve.

Also, a usual indicator is the Area under the ROC curve (AuROC), depicted in light blue. AuROC ranges between 0 and 1; 0.5 indicates a purely random detector and 1 is a perfect detector.

An equivalent quality indicator is 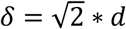, where *d* is the algebraic distance from the ROC curve to the first bisector. *δ* is defined by 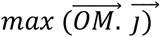, where M denotes an arbitrary point on the curve and 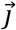 the vector (−1, 1). “ *δ* ” ranges between -1 and +1 (refer to Supplementary Information of this article). Therefore, for both AuROC and δ, higher values indicate a better method. As calculating and displaying the *δ* indicator is much less time consuming compared with AuROC, it was the chosen score for these networks and was termed ‘Distance to Diagonal ’.

##### Tests performed

The results given in this article (tables, figures) are derived from simulations.

#### 2.3 Networks with known dynamics

- Three-kinase network (three phosphorylated and bi-phosphorylated protein kinases: pRaF, ppMEK, ppERK), described in (Thomaseth *et al*., 2018). The ODEs related to the system dynamics are provided in the Supplementary Material of the quoted article. Perturbations applied to the total protein concentration: -80%, -10% and -1%.
- Linear three-gene network. Artificial network, composed of three genes with a linear dynamic behavior (created from (Alon, 2006)). Perturbations applied to the protein production rates: -80%, -10% and -1%.
- Four-node network, a schematic network of four genes, described in (Sontag *et al*., 2004). The ODEs are provided in the Supplementary Information of the quoted article (Supplementary Table 1). Perturbations applied to the maximum enzyme levels: -80%, -10% and -1%.
- MAPK cascade (6-node network), composed of three MAP kinases (MKKK, MKK and MAPK) and their bi-phosphorylated forms. Network described in (Kholodenko *et al*., 2002). ODEs are provided in the Supplementary Information of (Sontag *et al*., 2004) (Supplementary Table 2). Perturbations applied to the maximum enzyme levels: -80%, -50%, -10%, -1% and +50%.

These networks were digitally integrated from the ODEs, using the R “ode ” package. They were also used to compute the ANOVA described in § 1.3, with two technical replicates for each measurement (adding a slight Gaussian noise *N*(0, *σ*), *σ* =0.01 * mean of the non-perturbed expressions). For ease of reading, the ODEs are grouped in the Supplementary Information of the present article.

#### 2.4 Simulated networks

- Five 10-node networks, noted “InSilico_10_1 ” to “InSilico_10_5 ” and five 100-node networks (“InSilico_100_1 ” to “InSilico_100_5 ”), from DC4. Available from: https://www.synapse.org/#!Synapse:syn3049712/wiki/74630 Information about the corresponding data is in the Supplementary Information of the present article.
- Thirty networks generated by FRANK (Carré *et al*., 2017), corresponding to networks of 30, 60 and 100 nodes where all genes interact (TF = 30, 60 or 100 and TA = 0) and networks of 60, 100 and 200 nodes where half of the genes do not regulate any other gene (TF = TA = 30, 50 or 100). TF (Transcription Factor) is the number of nodes that regulate other nodes, and TA the number of nodes that regulate none. For each set, five networks were generated. The method used to generate these networks is described in (Borg *et al*., 2023) and is specified in the Supplementary Information of the present article.

In addition to the measurements of these thirty networks corresponding to the gene expression levels non-perturbed or perturbed by a KO (−100%) and a KD (−50%), noise was simulated by independent Gaussian random variables *N*(0, σ), σ = k * the mean concentration of genes. Two noise levels were simulated by giving to the coefficient of variation k the values 0.1 (medium noise) and 0.5 (strong noise). The solution, corresponding to the real connectivity coefficients, was also available (see Supplementary Information).

For all these networks (44), the exact solutions (connectivity matrices: “ *r*^*e*^ ”) were known and were used to compute the performance of the methods.

For the first four networks, generated from their ODE, and for the thirty FRANK networks, these solutions were obtained by calculation. These are matrices of real numbers. 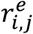 quantitatively represents the influence of node j on node i. To evaluate the studied developments with these networks, the Euclidean Distance was used (§ 2.1).

For the ten networks from DC4, the solution was provided by the DREAM Challenge website. These are Boolean matrices. 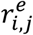 is 1 if node *j* influences node *i* and 0 otherwise. To evaluate the studied developments with these networks, the Distance to Diagonal (§ 2.2) was used.

### 3 “MRARegress ” package

The theoretical study allowed developing an easy-to-use software package, called MRARegress, that implements these advancements and checks the necessary conditions. This package, written in R, is freely available to the Scientific Community on GitHub (GNU General Public License v3.0 license), including the source code and the many unit tests data, covering 92% of the code (refer to Data availability statement).

This software requires R (version 4.3.2), RStudio (version ≥ 2023.06.0) and Cytoscape (version ≥ 3.6.1). Installation guidelines and simple examples for verification are provided in Supplementary Information as well as an overview of its main features. The list of R packages used and their version are indicated. A thumbnail presentation, an online manual and email support are available.

All results presented here were obtained using this software.

## Results and discussion

### 1 Statement of results

In a previous article (Borg *et al*., 2023), we demonstrated that MRA based on linear regression (MRARegress) substantially mitigates result estimation errors (connectivity coefficients) in the presence of noisy measurements or significant nonlinearity of the function (referred to as “ *f* ”) that describes the system dynamics. Here, we extended MRARegress:

- by permitting the use of not necessarily independent perturbations (given sufficient system rank),
- by calculating the 95% confidence intervals of the found connectivity coefficients,
- by enabling the detection of the estimation error primary source by ANOVA that takes into account the availability of replicates,
- by facilitating the use of polynomial regression (order 2) when error is primarily attributed to the nonlinearity of ‘ *f* ’ and noise levels are not excessive,
- by incorporating the user’s prior knowledge of the studied networks. We demonstrated this feasibility by seeking an optimal solution within a convex space.

### 2 Errors due to measurement noise

The error in estimating the coefficients 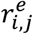 by the MRA method (||*r*^*e*^ − *r*||) originates from two sources:

- measurement noise (e.g., experimental conditions, measuring devices accuracy, operators ’ skills),
- the nonlinearity of function “ *f* ” in equation (2).

To decrease the measurement noise-induced error, it is advisable to increase the perturbation levels to boost the signal/noise ratio (Borg *et al*., 2023). Unfortunately, the perturbation level cannot always be chosen experimentally. In addition, increasing this level increases the nonlinearity-induced error. Hence, a balance must be found.

Alternatively, conventional methods involve multiplying measurements by providing multiple values to the *p*_*j*_ parameters, or performing successive measurements with the same parameter set (technical replicates), or on similar objects (e.g., several samples of the same cell line: biological replicates). While the classical MRA method only accommodates data averaging (Thomaseth *et al*., 2018), linear regression leverages over-determined systems (number of measurements exceeding N) far more effectively. In the presence of noisy measurements, this technique significantly reduces the result errors, as demonstrated in (Borg *et al*., 2023), (Santra *et al*., 2018).

### 3 Errors due to nonlinearity

Given the cost and difficulty of measurements, it is crucial to optimize the number of measurements and parameter values. One costly option entails employing the Design of Experiments theory that involves defining a plane of orthogonal experiments in the parameter space to minimize noise variance and improve the signal/noise ratio (Mason *et al*., 2003). The number of experiments required to infer the linear model connectivity coefficients is N.2^N-1^ for a N-node network, as opposed to at least N experiments with MRA.

The plethora of regression tools available facilitates a parsimonious approach concerning the number of experiments, on the order n_r_.N for linear models (n_r_: number of replicates per experiment).

Although the noise variance control is not optimal, the model quality can still be evaluated if the number of replicates is sufficient and experiments result in a strictly positive number of degrees of freedom (refer to § 1.3).

Analyzing the error variance (using ANOVA) allows identifying its primary cause and assessing whether the used linear model is appropriate (with the LOF test). If the error predominantly stems from the nonlinearity of function “ *f* ”, a polynomial regression (of order 2) should be used instead of linear regression.

We applied these methodologies to various networks (see Methods) in which three perturbations (−80%, -10%, and -1%) for the 3- and 4-node networks and five perturbations (−80%, -50%, -10%, -1%, and +50%) for the 6-node network were simulated. Each measurement had two replicates, corresponding to very low noise (independent Gaussian random variables of zero mean and standard deviation: 0.01*mean of non-perturbed expressions):

- 3-kinase network,
- linear 3-gene network,
- 4-node network,
- MAPK cascade (6 nodes).

Table 1 shows the results for the 3-kinase network.

**Table 1:** ANOVA results for the 3-kinase network.

| Variation | df | Sum of Squares | F-Value | p-Value |
| --- | --- | --- | --- | --- |
| Regression ($SSR$) | 2 | 10.08 ; 87.59 | 6312.53 ;<br>26831.38 | <1E-6 ; <1E-6 |
| Error ($SSE$) | 10 | 0.01 ; 0.05 | | |
| Lack of Fit error ($SSE_{LOF}$) | 4 | 0.01 ; 0.03 | 2.33 ; 25.87 | <1E-6; 0.17 |
| Pure error ($SSE_p$) | 6 | <1E-6; <1E-6 | | |
| Total | 12 |  |  |  |

(df: degrees of freedom; the value pairs, shown in the table and separated by semicolons, represent the min. and max. values for all nodes).

**Table 2:** ANOVA results for the Linear 3-gene network.

| Variation | df | Sum of Squares | F-Value | p-Value |
| --- | --- | --- | --- | --- |
| Regression ($SSR$) | 2 | 1.46 ; 5.51 | 485.54 ;<br>1629.26 | <1E-6 ; <1E-6 |
| Error ($SSE$) | 10 | 0.01 ; 0.02 | | |
| Lack of Fit error ($SSE_{LOF}$) | 4 | <1E-6 ; 0.01 | 0.04;3.94 | 0.07; 1 |
| Pure error ($SSE_p$) | 6 | <1E-6 ; 0.02 | | |
| Total | 12 |  |  |  |

The obtained results indicated that the p-value for *SSR* was <10^−6^. Consequently, the null hypothesis (H0) stating that “the value of Δ*X*_*i*_ is not correlated with the values of Δ*X*_*k*_, *k*≠*i* ” could be rejected for all nodes.

Conversely, the p-value for *SSE*_*LOF*_ ranged between 0 and 17%. For some nodes, the null hypothesis “residual errors stem from measurement noise ” could be rejected with 95% certainty, indicating that for these nodes, error originated mostly from the chosen linear model, rendering linear regression inadequate. This concerned two of the three nodes according to the MRARegress software.

In the linear 3-gene network, the null hypothesis “residual errors stem from measurement noise ” could not be rejected with 95% certainty for any node, consistent with the linear system of equations that described the dynamics (*SSE*_*LOF*_ p-value ≥ 7% for all nodes).

For the 4-node and MAPK cascade networks, results were similar to those obtained for the 3-kinase network. Two of the four nodes (4-node network) and five of the six nodes (MAPK cascade) displayed p-values for *SSE*_*LOF*_ lower than the 5% threshold, indicating once more that most error stemmed from the selected model, thus rendering the linear regression unsuitable.

The detailed results for each network (F-value and p-value for each node) are in Supplementary Information.

Figure 5 describes the discovered networks using the specified perturbations with two replicates.

**Figure 5:**
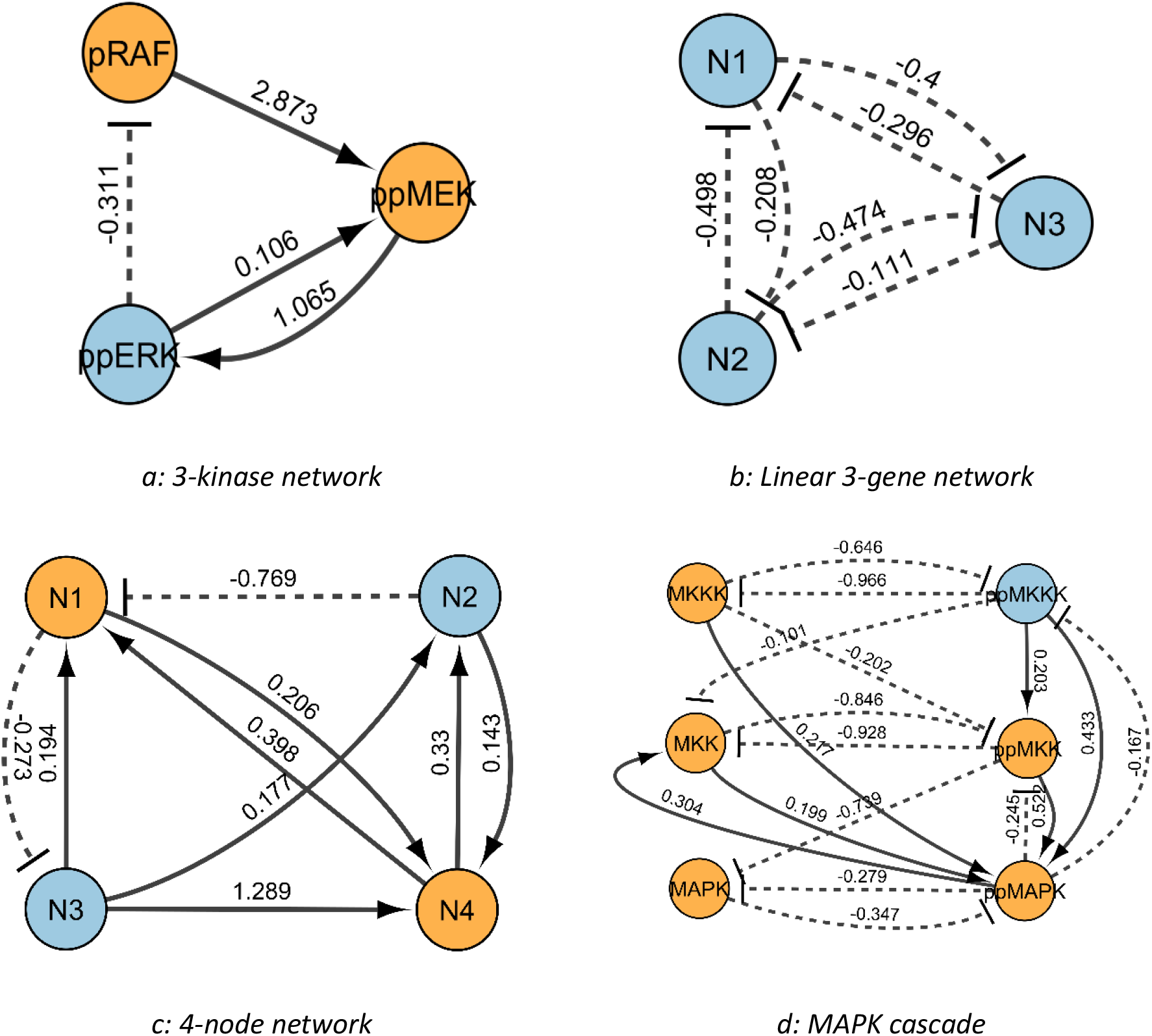
Illustration of non-linearity highlighted by MRARegress. In orange, nodes for which linear regression was unsuitable. The values associated with the edges were the calculated connectivity coefficients r_i,j_.

### 4 Polynomial regression (order 2)

ANOVA indicated the presence of significant error due to nonlinearity for several nodes within these networks. To evaluate the utility of processing this nonlinearity, we computed the L2-norm distance between the exact connectivity matrix 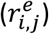 and the approximate connectivity matrix (*r*_*i,j*_), calculated through the prescribed perturbations, for the three nonlinear networks (3-kinase, 4-node and MAPK cascade) using the linear MRARegress and polynomial MRARegress (second degree) methods.

Results are summarized in Table 3 and the detailed matrices are provided in Supplementary Information.

**Table 3:** Euclidean distance between exact and approximate connectivity matrices (noise-free measurements)

| Network | 3 kinases | 4 genes (4 nodes) | MAPK cascade (6 nodes) |
| --- | --- | --- | --- |
| Perturbations | -80%, -10%, -1% | -80%, -10%, -1% | -80%, -50%, -10%, -1%, +50% |
| Distance (Linear MRA) | 0.254 | 0.617 | 0.871 |
| Distance (2<sup>nd</sup> degree MRA) | 0.009 | 0.002 | 0.036 |

The superiority of employing second-order polynomial regression is evident, although this method introduces more unknowns than linear regression, rendering results more susceptible to measurement noise.

We performed the sensitivity analysis by simulating measurement noise at different levels (Gaussian noise *N*(0, *σ*), *σ* = k * mean of the non-perturbed network node expression level; see Methods). This demonstrated the advantage of using ANOVA for gene expression measurements. If ANOVA reveals that the primary source of errors stems from the function *f* nonlinearity and measurement noise is low, employing second-order polynomial regression is highly advantageous; otherwise, linear regression remains preferable.

The obtained results are listed in Table 4 (average over 20 simulations for each value of the variation coefficient k). Networks, perturbations and replicates were the same as in Table 3.

**Table 4:**
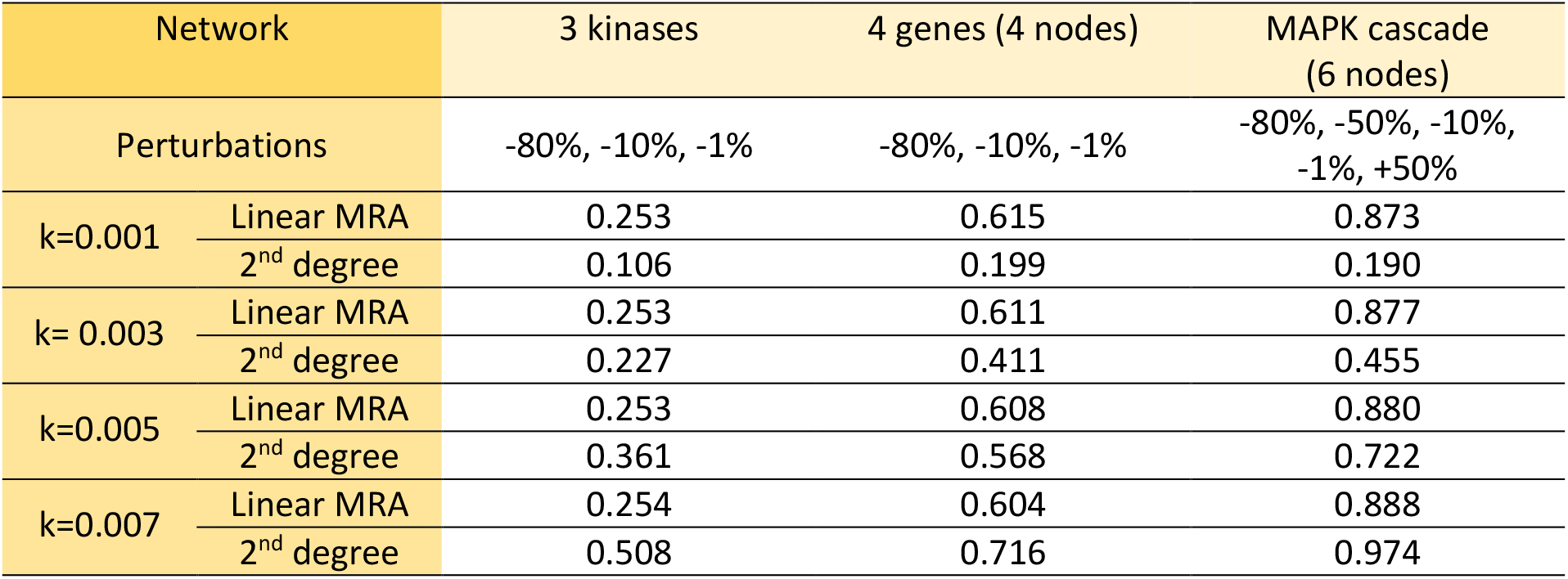
Euclidean distances between exact and approximate connectivity matrices, depending on the noise level.

| Network | 3 kinases | 4 genes (4 nodes) | MAPK cascade (6 nodes) |
| --- | --- | --- | --- |
| Perturbations | -80%, -10%, -1% | -80%, -10%, -1% | -80%, -50%, -10%, -1%, +50% |
| k=0.001 Linear MRA | 0.253 | 0.615 | 0.873 |
| k=0.001 2<sup>nd</sup> degree | 0.106 | 0.199 | 0.190 |
| k= 0.003 Linear MRA | 0.253 | 0.611 | 0.877 |
| k= 0.003 2<sup>nd</sup> degree | 0.227 | 0.411 | 0.455 |
| k=0.005 Linear MRA | 0.253 | 0.608 | 0.880 |
| k=0.005 2<sup>nd</sup> degree | 0.361 | 0.568 | 0.722 |
| k=0.007 Linear MRA | 0.254 | 0.604 | 0.888 |
| k=0.007 2<sup>nd</sup> degree | 0.508 | 0.716 | 0.974 |

### 5 Contribution of prior knowledge

Linear regression offers many advantages to the MRA method due to extensive mathematical developments that facilitate module network inference. For example, various cost functions can be used to consider the network characteristics (e.g., size, sparsity) and the importance attributed to specific parameters. Different methods corresponding to these cost functions (e.g., LSE, LASSO, RIDGE, Elastic Net, STEP) have been compared (Borg *et al*., 2023) for problem dimension reduction and for eliminating irrelevant network edges.

Whatever the cost function, the MRARegress method typically involves solving *N* optimization problems with values in ℝ^*N*−1^. Indeed, generally nothing is known about the *N-1* connectivity coefficients *r*_*i,j*_ (j ∈ ⟦1, *N*⟧, *j*≠*i*).

However, in some cases, prior information on these values may be available (e.g., biological knowledge, databases), such as:

- the *j* module has no effect on the *i* module (therefore, *r*_*i,j*_ = 0), or
- the *j* module amplifies the action of the *i* module (therefore, *r*_*i,j*_ ≥ *α*, where *α* ≥ 0)
- the *j* module inhibits the action of the *i* module (therefore, *r*_*i,j*_ ≤ −*α*, where *α* ≥ 0) or
- the coefficient *r*_*i,j*_ has a value between *a* and *b* (therefore *a* ≤ *r*_*i,j*_ ≤ *b*), or
- there is a linear relationship between specific coefficients *r*_*i,j*_ etc …

The incorporation of this information implies that the solutions of the optimization problems are not in ℝ^*N*−1^, but lie within a convex subspace of it.

*Reminder: A* C ℝ^*N*^*is a convex subspace of* ℝ^*N*^, *if*

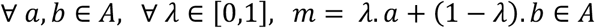

Furthermore, leveraging regression-based methods grants access to a comprehensive mathematical toolkit. For instance, the CVXR library of R functions, developed by João Neto based on the work by (Boyd and Vandenberghe, 2004), addresses convex optimization problems (http://www.di.fc.ul.pt/~jpn/r/optimization/CVXR.html).

This library is applicable when the cost function used and all inequality type constraints on the unknown *r*_*i,j*_ are convex functions, and all equality type constraints are affine. This suitability can be met by using the least squares method as a cost function (also the Threshold Linear Regression method, called TLR, of the MRARegress package, see Supplementary Information).

*Reminder: f, A C* ℝ^*N*^→ ℝ *is a convex function if A is a non-empty convex subspace of* ℝ^*N*^*and if*

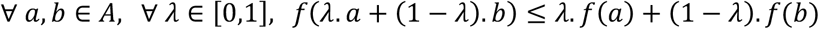

The efficacy of this knowledge in terms of performance was assessed using two test files (25 networks in total). These files contained measurement data (networks perturbed by a KO and a KD of 50%, and non-perturbed networks) and exact solutions:

- Five 10-node networks and five 100-node networks from DC4. Measurements were noisy, of unknown type and level. The provided exact solution only indicated the existence or absence of an edge between nodes (i.e.,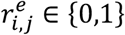).
- Five networks of 30, 60 and 100 nodes, composed of genes that regulate at least another gene, generated by FRANK (Carré *et al*., 2017). Undisturbed and perturbed (KO, KD) data, with two noise levels, were available as well as the exact solution 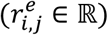.

We compared the simulation results with the known solutions.

To simulate a designated percentage “*p* ” of knowledge, we randomly chose (uniform distribution) a set of pairs (*i, j*), where *i* ∈ ⟦1, *N*⟧, *j* ∈ ⟦1, *N*⟧, *i*≠*j*. The number of pairs was 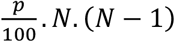, rounded to the nearest integer. The number of unknown elements in the *(N, N)* connectivity matrix “ r ” was *N*(N-1)* because the elements of the diagonal were -1. For each selected pair, we specified the action of module *j* on module *i* (amplification, inhibition, no action) to the program.

Then, we computed the approximate connectivity matrix (*r*) using the MRARegress method. For networks resulting from DC4, this matrix was digitized to obtain a matrix named “*rdig* ”, via the TLR method introduced in (Borg *et al*., 2023) and described in Supplementary Information.

We determined the scores by calculating the Distance to Diagonal (DC4 networks; Fig. 6) or the Euclidean distance (FRANK-generated networks; Fig. 7) between the matrices “ *r* ” and “ *r*^*e*^ ”. For each studied network and percentage value, ten random draws of pairs *(i,j*) were conducted. The mean and standard deviation of the obtained scores were computed across these ten draws. The selected score corresponded to this mean.

**Figure 6:**
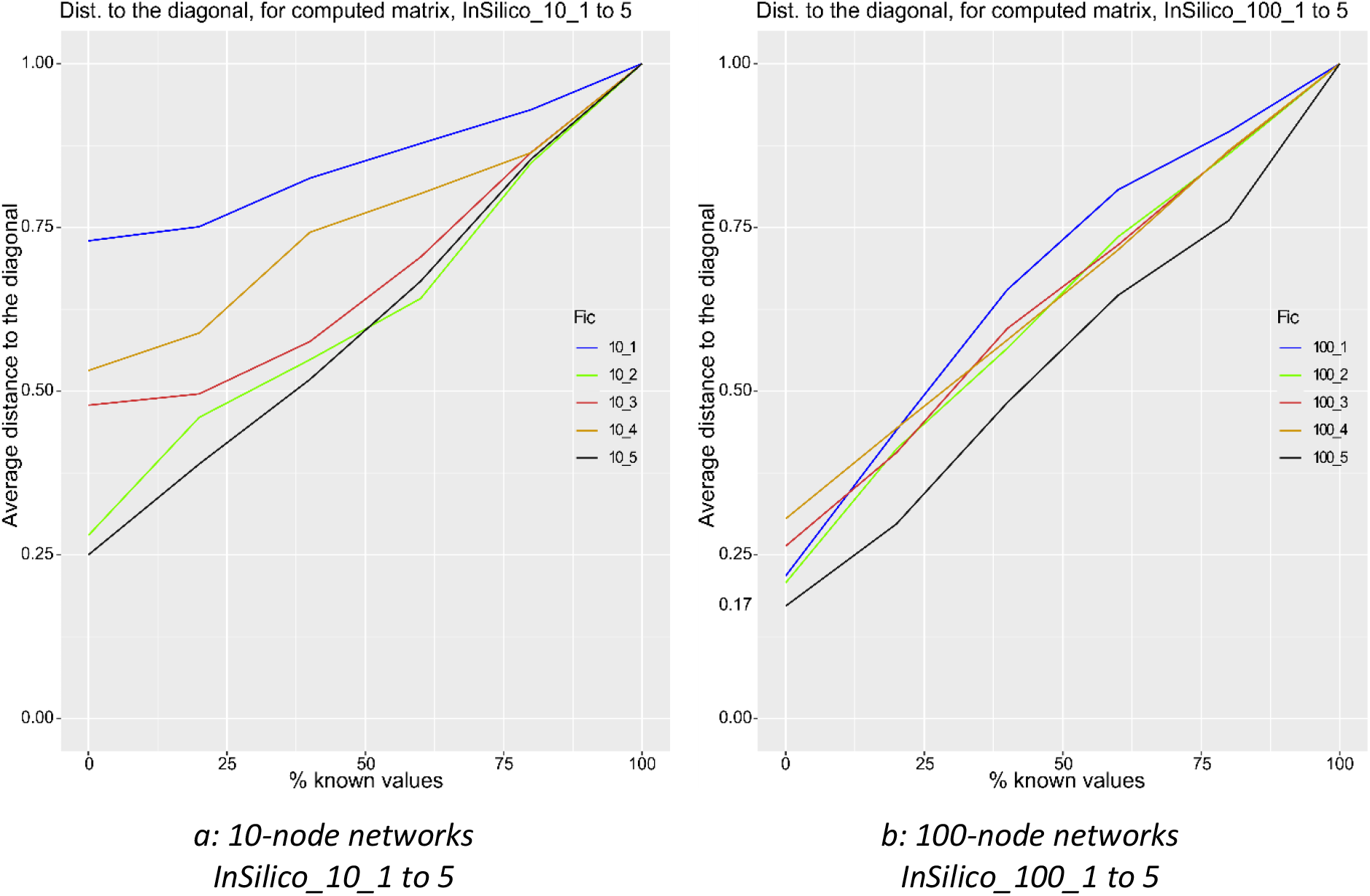
Mean values of the Distance to Diagonal of the DC4 networks based on the percentage of known data (10 random draws for each percentage). To maintain readability, the standard deviations are not displayed in the figures, but are provided in Supplementary Information.

**Figure 7:**
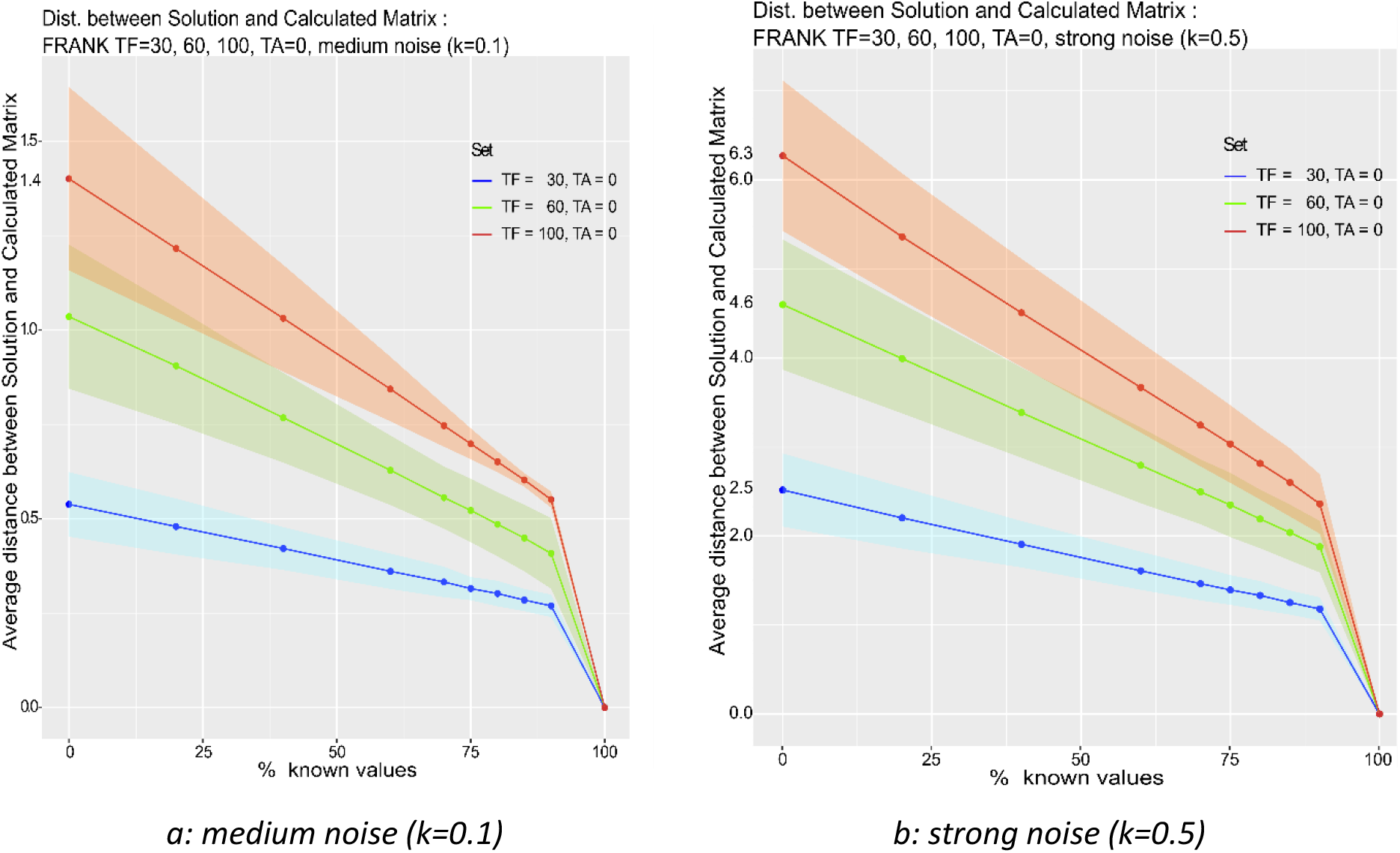
Average distance between solution and calculated matrix for FRANK-generated networks (30, 60, 100 nodes) in function of the percentage of known data and noise level. Each curve shows the mean value ± standard deviation (smoothing line) – 5 networks for each curve.

Figure 6 shows these scores, in function of “*p* ”, for the five 10-node networks (Fig. 6a) and five 100-node networks (Fig. 6b), from DC4.

Figure 7 shows the mean of the scores (Euclidean distance) in function of “*p* ” for the three sets of five networks of 30, 60 and 100 nodes generated by FRANK (TF = 30, 60 or 100 and TA = 0) at two noise levels (medium: k = 0.1 and strong: k = 0.5, as described in Methods).

These results underscore the importance of incorporating prior knowledge and the consequent substantial performance enhancement. Additionally, the very small standard deviation values indicated that this improvement predominantly relies on the percentage of known values rather than on their specific location.

To complement this study, we used again the FRANK Network Generator to simulate three networks with 30, 50 and 100 genes where each gene regulated at least another gene and an equivalent number of genes regulated none (out degree = 0). These networks (60, 100 and 200 nodes) were subject to the same perturbations and noise levels as before. The exact solution was known. To simulate a specified percentage “*p* ” of knowledge, we randomly selected (uniform distribution) 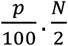 integers that represented the corresponding non-regulating node numbers and communicated them to the program (*N*∈ {60,100,200}). The score was calculated as the Euclidean distance between the exact and computed connectivity matrices.

Figure 8 illustrates the mean scores ± standard deviation (smoothing line) corresponding to the ten draws, in function of “*p* ”, for the sets of five networks with 60, 100,and 200 nodes.

**Figure 8:**
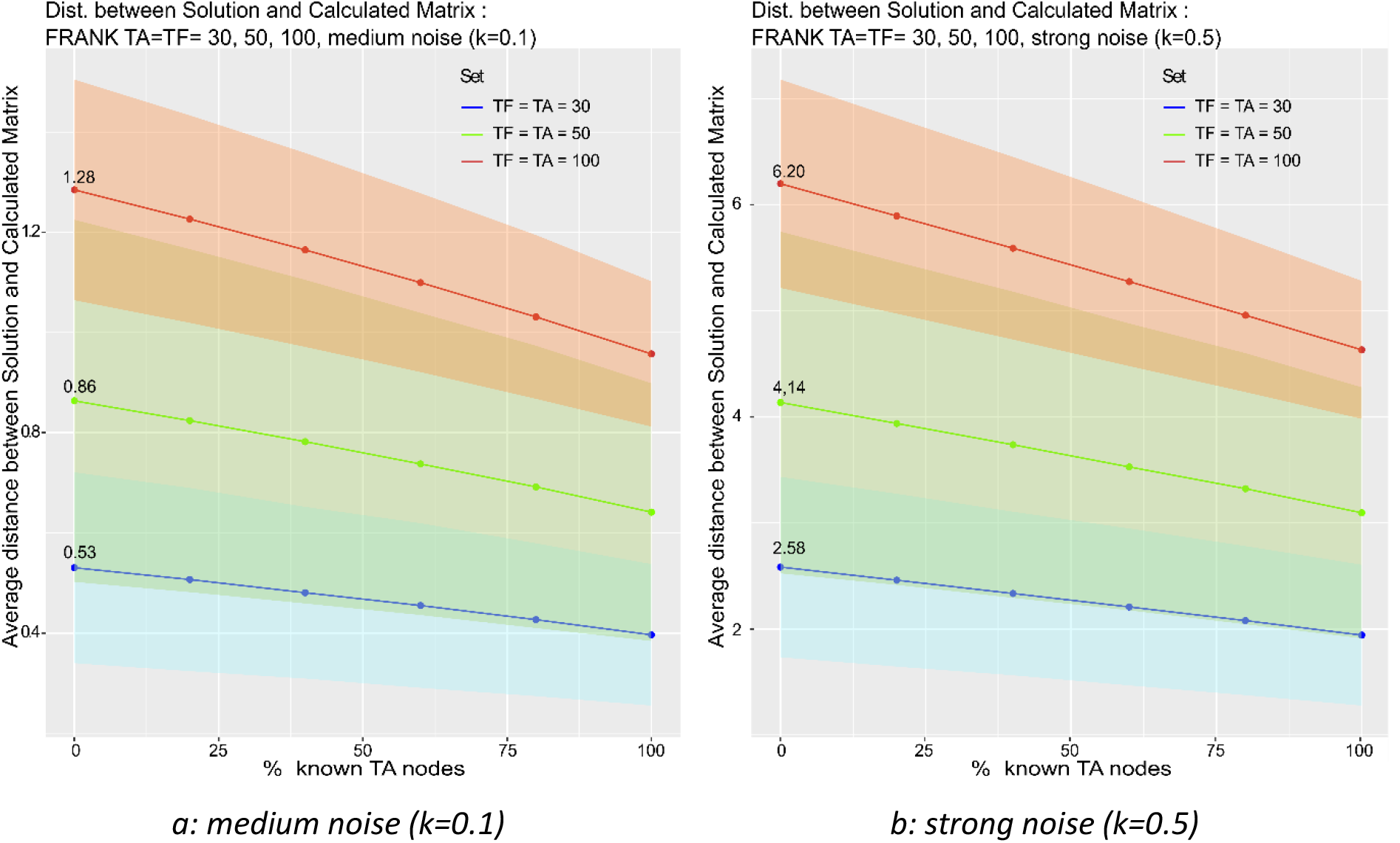
Average distance between solution and calculated matrix for FRANK-generated networks (60, 100, 200 nodes, including 50% of non-regulating genes) based on the percentage of known non-regulating genes and noise level (mean ± standard deviation – 5 networks for each curve).

## Conclusion

Understanding biological systems often involves the meticulous inference of interaction networks. This prompted the development of various methods, including MRARegress. MRARegress was introduced to efficiently address inference in the presence of noise, which is inevitable, and for networks larger than 10 nodes. In this study, we described and validated strategies to overcome some limitations of MRA, such as the prerequisite of perturbation-independence or the near-linearity of the function that describes the system dynamics. We also highlighted the usefulness of polynomial regression in conditions of low noise and nonlinear behavior and the substantial benefit provided by prior knowledge about the studied network, when available. All these advancements have been integrated into our MRARegress software package. This package allows:

- the application of different methods (e.g., ARACNE, LSE, TLR, LASSO, STEP, Random Forest) to analyze measurements
- the optimal consideration of over-determined systems and replicates,
- the accommodation of non-independent perturbations (subject to system rank) to incorporate different experiment designs,
- the calculation of the 95% confidence interval for the computed connectivity coefficients,
- ANOVA analysis to identify nodes that are unsuitable for linear regression and to what extent,
- the use of polynomial regression (order 2) to compute synergy-related coefficients alongside connectivity coefficients,
- the incorporation of any prior knowledge on the network under study.

The research landscape surrounding MRA continues to evolve. Future endeavors may include automating program hyper-parameters, integrating additional artificial intelligence-based processing functionalities, preprocessing measurements based on their noise characteristics, and analyzing networks with periodic or pseudo-periodic behavior while considering the measurement changes over time. MRARegress is an open-source software and this should allow integrating these potential advancements.

### Key Points

- MRA, integrated with linear regression (“MRARegress ”), is an effective method for inferring biological networks, including noisy and large networks.
- Regression helps to overcome some MRA limitations, such as perturbation independence. The ANOVA and LOF tests identify the primary source of errors (measurement noise or nonlinearity). If nonlinearity prevails, a second order polynomial regression significantly improves the results.
- MRARegress facilitates the use of prior knowledge on the studied network. It allows reducing almost linearly the estimation error of network connectivity coefficients.
- These accomplishments are implemented in a software package, also named “MRARegress ”, developed in R and freely available to the scientific community.

## Data availability statement

The source code and the unit tests data are available online at https://github.com/J-P-BORG/MRARegress

The files containing all calls to MRARegress and its modules and the corresponding parameters (AdvancedMRA.R) and all the data files used to obtain the results in this article (tables, figures) are also available on line at https://github.com/J-P-BORG/MRA or https://github.com/J-P-BORG/MRARegress, within the data folder.

## Notes

### Competing Interest Statement

The authors have declared no competing interest.

