## Supplementary material for "Testing and overcoming the limitations of Modular Response Analysis": Suplementary information

### 1 "TLR" method and "Distance to Diagonal"

The" Threshold Linear Regression" (TLR) method involves:

1. Employing the MRARegress algorithm to search for the connectivity matrix, utilizing the Least Square Error (LSE) as the cost function.
2. Identifying the maximum absolute values of connectivity coefficients.
3. Setting connectivity coefficients $r_{i,j}$ to 0 if their absolute value is less than or equal to the threshold $0.25*max(\left| r_{i,j} \right|)$..

In employing this method to digitize a connectivity matrix, coefficients with absolute values exceeding the threshold are set to 1.

The selection of coefficient 0.25 was determined through extensive testing on simulated networks. Apart from its efficacy, the method's simplicity ensures computational efficiency by solely requiring the calculation of a threshold value. This digitization approach is instrumental in computing the "distance to diagonal" (δ), with the threshold value corresponding to the $M_{0}$ point.

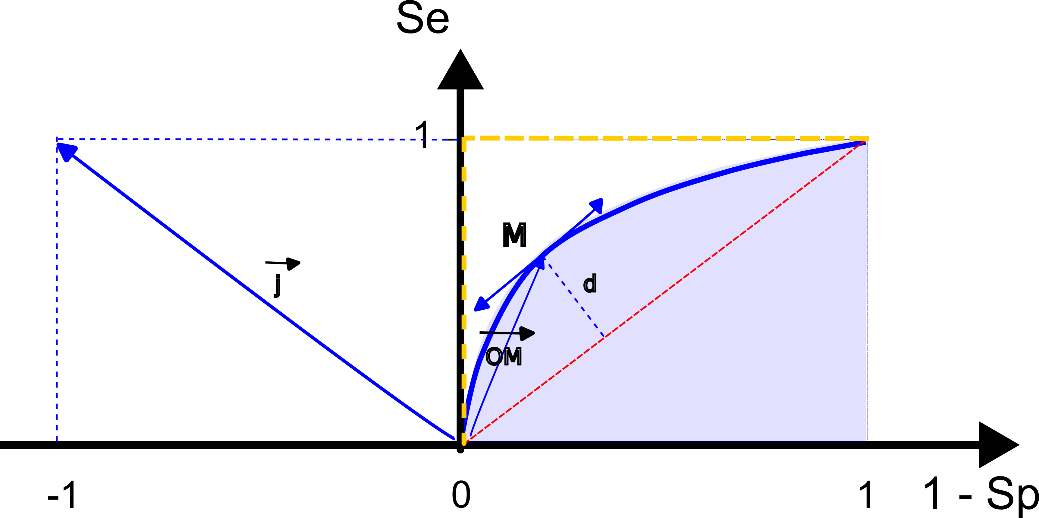

"Distance to Diagonal" $\delta$ is defined by $max (\vec{OM.} \vec{j),}$ where M denotes an arbitrary point on the curve, coordinates of O are (0,0) and $\vec{j} is$the vector (-1, 1). We can check that $\left\| \delta\right\|= \sqrt{2}*d$, so "$\delta$" ranges between -1 and +1.

Replacing $\delta=max(\vec{OM.} \vec{j})$ with $\delta=\vec{{OM}_{0}}. \vec{j}$, introduces a minor error. However, this error, though slight, is directionally correct, as the obtained results tend to be marginally lower than those derived from $\max\left( \vec{OM.} \vec{j} \right)$ calculations. Moreover, since this method is uniformly applied across cases where distance to the diagonal is utilized, it doesn't introduce bias when comparing results.

Furthermore, computational time is significantly reduced, as only one single scalar product calculation is required, as opposed to an integral calculation in the AuROC scenario.

### 2 Networks with known dynamics

For all aforementioned networks, exact connectivity matrix data were computed by

$${\forall k\neq i, r}_{i,k}^{e}=\frac{\partial\varphi_{i}}{\partial X_{k}}\left( P0 \right)= -\frac{\frac{\partial f_{i}}{\partial X_{k}}}{\frac{\partial f_{i}}{\partial X_{i}}}\left( P0 \right) and r_{i,i}^{e}=-1.$$

While the intricate calculations are omitted here, the results are retrievable from specified files indicated in the following paragraphs. Approximating these results is feasible by subjecting noise-free networks to minimal perturbations.

#### 2.1 "3 kinases" network

##### 2.1.1 Equations of dynamics

Fom C. Tomaseth et al's "Supplementary Material" S1 a (Thomaseth *et al.*, 2018)

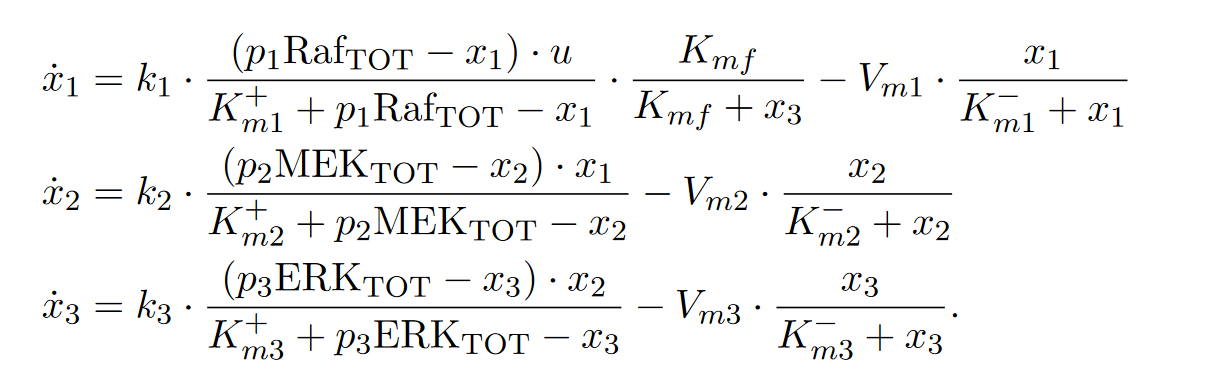

where: p1, p2, p3 are multiplicative coefficients, representing the total amount of protein and perturbations performed.

k_1_ = 5, k_2_ = 3, k_3_ = 1, u = 1,

Raf_TOT_ = MEK_TOT_ = ERK_TOT_ = 20,

$K_{m1}^{+}= K_{m2}^{+}= K_{m3}^{+}= K_{m1}^{-}= K_{m2}^{-}= K_{m3}^{-}=$20, K_mf_ = 5,

V_m1_ = V_m2_ = V_m3_ = 10 and

x1(t), x2(t), x3(t)] refer to the active states of the three proteins pRaf, ppMEK and ppERK.

##### 2.1.2 ANOVA

| Node | SSR | | | | LOF | | | |
| --- | --- | --- | --- | --- | --- | --- | --- | --- |
|  | df | Σ squares | F | pValue | df | Σ squares | F | pValue |
| pRAF | 2 | 10.08 | 8084.2 | 1E-16 | 4 | 0.005 | 8.37 | 0.01 |
| ppMEK | 2 | 65.64 | 6312.5 | 3E-16 | 4 | 0.034 | 25.87 | 0.00 |
| ppERK | 2 | 87.59 | 26831.4 | 2E-19 | 4 | 0.008 | 2.33 | 0.17 |

##### 2.1.3 Exact and approximate connectivity matrices

| $\left( \begin{matrix} -1 & 0 & -0.18 \\ 3.02 & -1 & 0 \\ 0 & 0.98 & -1 \end{matrix} \right)$  a : Exact Connectivity Matrix | $\left( \begin{matrix} -1 & 0 & -0.31 \\ 2.85 & -1 & 0.11 \\ -0.01 & 1.07 & -1 \end{matrix} \right)$  b : Approximate Connectivity Matrix (using order 1) | $\left( \begin{matrix} -1 & 0 & -0.18 \\ 3.03 & -1 & 0 \\ 0.01 & 0.98 & -1 \end{matrix} \right)$  c : Approximate Connectivity Matrix (using order 2) |
| --- | --- | --- |

Exact connectivity matrix : file "Solution_3G.rda".

d ($r^{e},r)$ = 0.2535509	Order 1

d ($r^{e},r)$ = 0.00876365	Order 2

#### 2.2 "Linear 3 genes" network

Following Alon's suggestions (Alon, 2006), a purely artificial network comprised of three genes exhibiting linear dynamic behavior was simulated to evaluate "Lack Of Fit" under controlled conditions.

Gene 1 regulates protein kinase production, pivotal in phosphorylating other proteins.

Gene 2 regulates phosphatase protein production, facilitating the dephosphorylation of proteins phosphorylated by Gene 1's protein kinase.

Gene 3 regulates regulatory protein production, involved in signaling pathway protein interactions.

Arbitrary yet realistic constants were chosen for this simulation.

##### 2.2.1 Equations of dynamics

Possible parameters:

• Protein production rate:

- α1=0.5	(unit of concentration per unit of time)
- α2=0.4
- α3=0.3

• Protein modification rate:

- β1=0.1	(inverse time unit: 1/t)
- β2=0.2
- β3=0.15

$$\left\{ \begin{aligned} \frac{dx_{1}}{dt}=0.5-0.1*x_{1}-0.05*x_{2}-0.03*x_{3} \\ \frac{dx_{2}}{dt}=0.4-0.04*x_{1}-0.2*x_{2}-0.02*x_{3} \\ \frac{dx_{3}}{dt}=0.3-0.06*x_{1}-0.07*x_{2}-0.15*x_{3} \end{aligned} \right.$$

##### 2.2.2 ANOVA

| Node | SSR | | | | LOF | | | |
| --- | --- | --- | --- | --- | --- | --- | --- | --- |
|  | df | Σ squares | F | pValue | df | Σ squares | F | pValue |
| N1 | 2 | 1.65 | 673.92 | 2E-11 | 4 | 0.008 | 3.94 | 0.067 |
| N2 | 2 | 1.46 | 485.54 | 1E-10 | 4 | 0.009 | 1.99 | 0.216 |
| N3 | 2 | 5.51 | 1629.26 | 3E-13 | 4 | 0.000 | 0.035 | 0.997 |

##### 2.2.3 Exact and approximate connectivity matrices

| $\left( \begin{matrix} -1 & -0.5 & -0.3 \\ -0.2 & -1 & -0.1 \\ -0.4 & -0.47 & -1 \end{matrix} \right)$  a : Exact Connectivity Matrix | $\left( \begin{matrix} -1 & -0.5 & -0.3 \\ -0.21 & -1 & -0.11 \\ -0.4 & -0.47 & -1 \end{matrix} \right)$  b : Approximate Connectivity Matrix (using order 1) |
| --- | --- |

Exact connectivity matrix : file "Solution_3L.rda".

For this linear network, an approximate connectivity matrix of order 1 was constructed using marginally noisy data and two replicates. As noise-free data perfectly aligns, the "lm" program issuing the linear regression provides a warning: "Near perfect fit: summary may not be reliable", hence refraining from calculation.

The LOF test conducted via ANOVA in § 2.2.2 confirms the acceptability of the linear model for all three nodes, obviating the need for approximating a second-order connectivity matrix.

#### 2.3 "4 nodes" network

##### 2.3.1 Equations of dynamics

From Sontag et al's "Supplementary Information" : Supplementary Table 1 copied here for convenience (Sontag *et al.*, 2004).

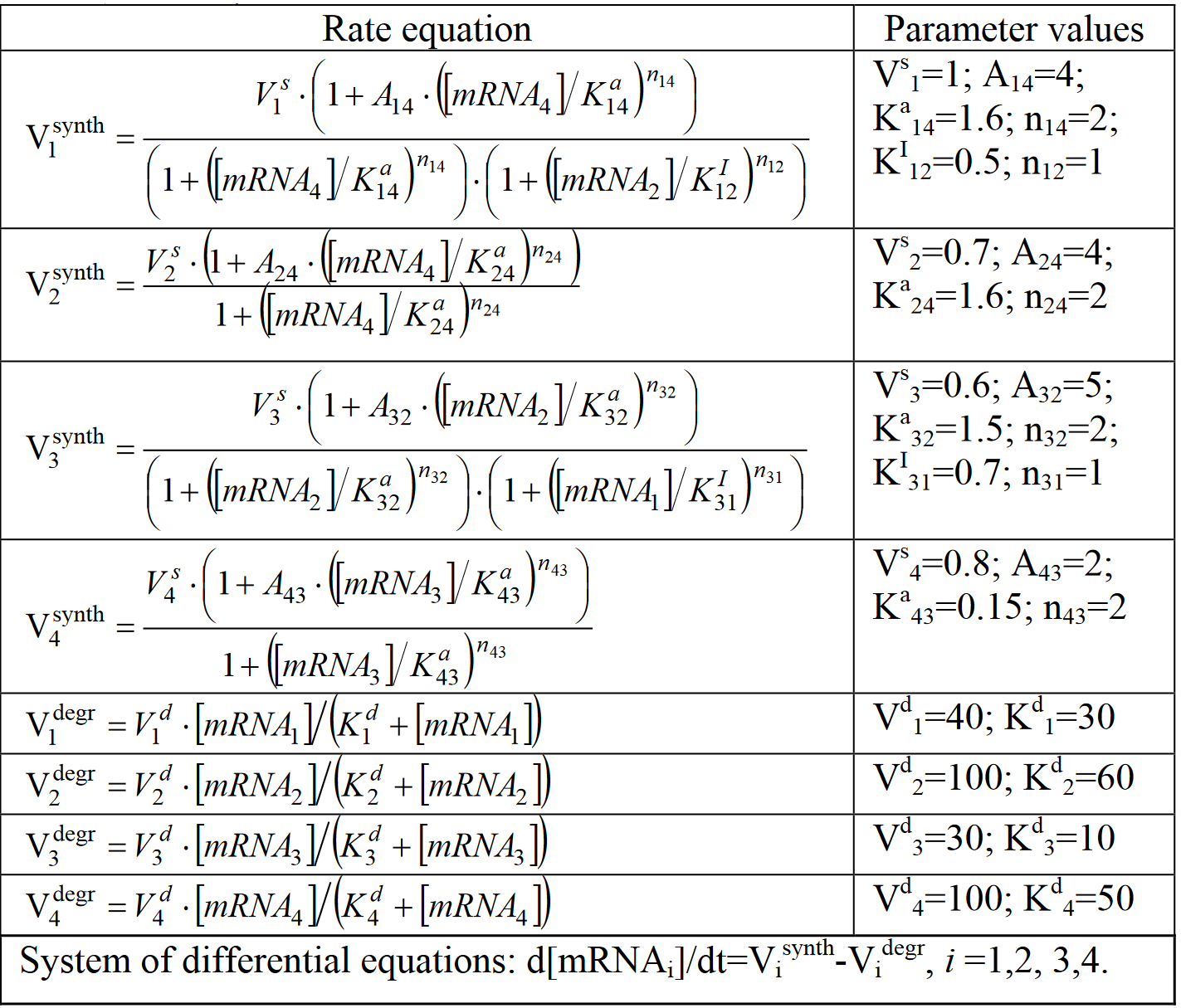

The parameters serve as multiplicative coefficients for maximum enzyme levels $V_{1}^{s} to V_{4}^{s}.$

##### 2.3.2 ANOVA

| Node | SSR | | | | LOF | | | |
| --- | --- | --- | --- | --- | --- | --- | --- | --- |
|  | df | Σ squares | F | pValue | df | Σ squares | F | pValue |
| N1 | 3 | 0.22 | 970.82 | 2E-17 | 6 | 0.001 | 9.58 | 0.002 |
| N2 | 3 | 0.09 | 511.20 | 2E-15 | 6 | 0.000 | 2.95 | 0.071 |
| N3 | 3 | 0.04 | 485.22 | 4E-15 | 6 | 0.000 | 2.01 | 0.166 |
| N4 | 3 | 0.14 | 499.67 | 3E-15 | 6 | 0.001 | 9.60 | 0.002 |

##### 2.3.3 Exact and approximate connectivity matrices

| $\left( \begin{matrix} -1 & -0.45 & 0 & 0.39 \\ 0 & -1 & 0 & 0.48 \\ -0.16 & 0.19 & -1 & 0 \\ 0 & 0 & 1.05 & -1 \end{matrix} \right)$  a : Exact Connectivity Matrix | $\left( \begin{matrix} -1 & -0.78 & 0.17 & 0.40 \\ 0.01 & -1 & 0.16 & 0.33 \\ -0.25 & 0 & -1 & 0.07 \\ 0.18 & 0.13 & 1.36 & -1 \end{matrix} \right)$  b : Approximate Connectivity Matrix (using order 1) | $\left( \begin{matrix} -1 & -0.45 & 0 & 0.39 \\ 0 & -1 & 0 & 0.48 \\ -0.16 & 0.19 & -1 & 0 \\ 0 & 0 & 1.05 & -1 \end{matrix} \right)$  c : Approximate Connectivity Matrix (using order 2) |
| --- | --- | --- |

Exact connectivity matrix : file "Solution_4G.rda".

d ($r^{e},r)$ = 0.6169653	Order 1

d ($r^{e},r)$ = 0.002214955	Order 2

#### 2.4 "MAPK Cascade" (6 nodes)

##### 2.4.1 Equations of dynamics

From Sontag et al's "Supplementary Information" : Supplementary Table 2 copied here for convenience (Sontag *et al.*, 2004).

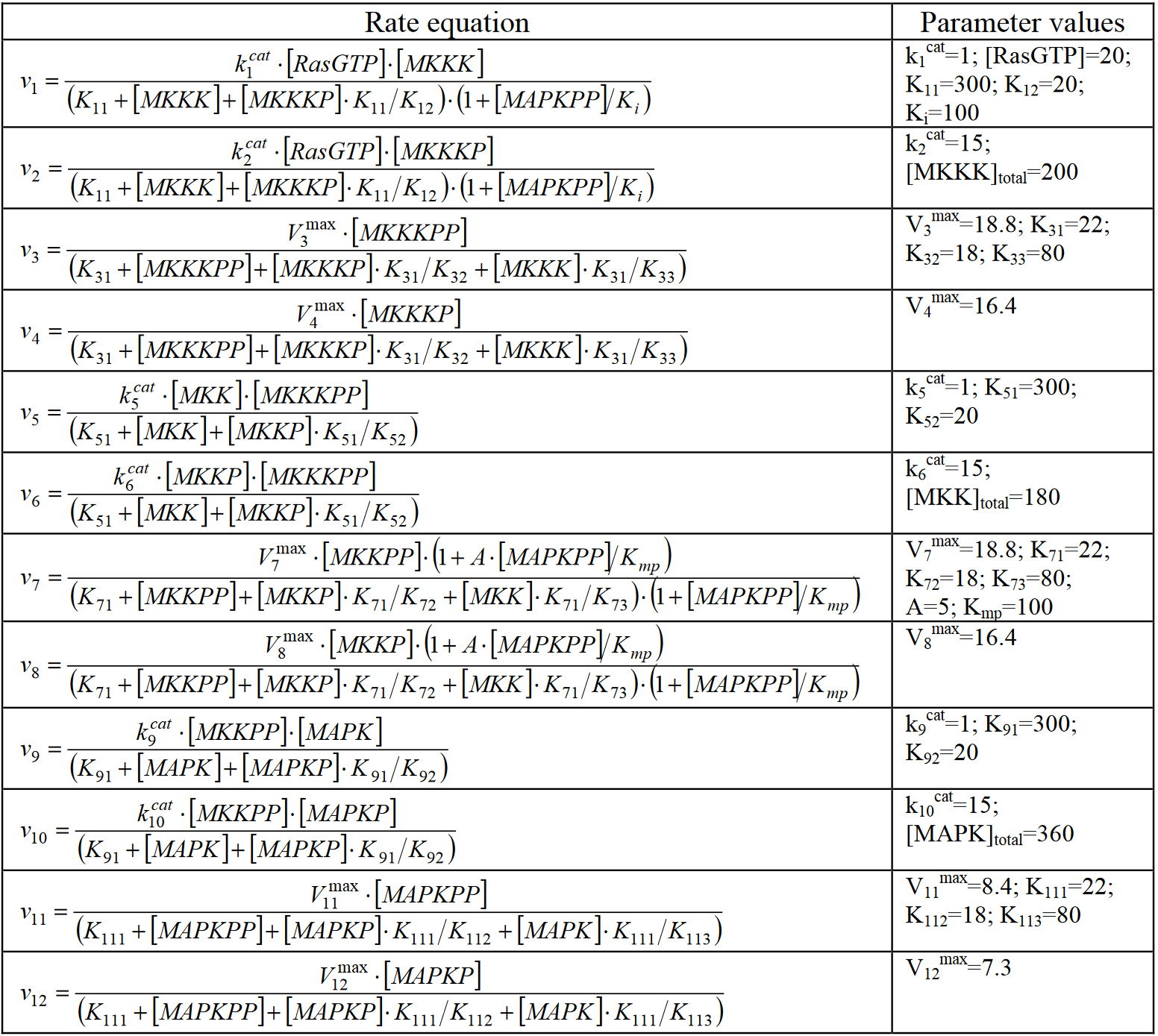

The kinetic equations and moiety conservations derived from the stoichiometry are the following:

d[MKKK-P]/dt = *v*_1_ - *v*_2_ + *v*_3_ – *v*_4_;

d[MKKK-PP]/dt = *v*_2_-*v*_3_;

d[MKK-P]/dt = *v*_5_ – *v*_6_ + *v*_7_ – *v*_8_;

d[MKK-PP]/dt = *v*_6_ – *v*_7_;

d[MAPK-P]/dt = *v*_9_ – *v*_10_ + *v*_11_ – *v*_12_;

d[MAPK-PP]/dt = *v*_10_ – *v*_11_;

[MKKK]_total_ = [MKKK] + [MKKK-P] + [MKKK-PP];

[MKK]_total_ = [MKK] + [MKK-P] + [MKK-PP];

[MAPK]_total_ = [MAPK] + [MAPK-P] + [MAPK-PP].

By inserting the conservation equations in the differential equations, we obtained a system of six differential equations that was integrated numerically with the "ode" R package

##### 2.4.2 ANOVA

| Nodes | SSR | | | | LOF | | | |
| --- | --- | --- | --- | --- | --- | --- | --- | --- |
|  | df | Σ  squares | F | pValue | df | Σ squares | F | pValue |
| MKKK | 5 | 21452.08 | 1129.30 | 4E-46 | 20 | 75.93 | 2.38 | 0.021 |
| ppMKKK | 5 | 4165.21 | 331.71 | 3E-34 | 20 | 43.67 | 0.98 | 0.517 |
| MKK | 5 | 27258.43 | 1192.29 | 1E-46 | 20 | 123.78 | 4.23 | 4E-4 |
| ppMKK | 5 | 29528.79 | 870.82 | 1E-43 | 20 | 222.77 | 8.43 | 1E-6 |
| MAPK | 5 | 28960.16 | 1077.92 | 1E-45 | 20 | 191.45 | 6.45 | 1E-5 |
| ppMAPK | 5 | 18474.49 | 310.77 | 1E-33 | 20 | 436.48 | 9.78 | 2E-7 |

##### 2.4.3 Exact and approximate connectivity matrices

| $\left( \begin{matrix} -1 & -0.99 & 0 & 0 & 0 & 0.12 \\ -0.71 & -1 & 0 & 0 & 0 & -0.16 \\ 0 & -0.26 & -1 & -0.85 & 0 & 0.16 \\ 0 & 0.59 & -0.89 & -1 & 0 & -0.35 \\ 0 & 0 & 0 & -0.71 & -1 & -0.66 \\ 0 & 0 & 0 & 0.27 & -0.28 & -1 \end{matrix} \right)$  a : Exact Connectivity Matrix |  |
| --- | --- |
| $\left( \begin{matrix} -1 & -0.97 & 0 & -0.01 & 0 & 0.12 \\ -0.64 & -1 & 0.02 & 0.03 & 0 & -0.16 \\ 0.05 & -0.06 & -1 & -0.92 & 0.01 & 0.24 \\ -0.17 & 0.29 & -0.85 & -1 & 0 & -0.33 \\ 0.02 & 0.05 & 0.10 & -0.75 & -1 & -0.30 \\ 0.25 & 0.44 & 0.26 & 0.58 & -0.36 & -1 \end{matrix} \right)$  b : Approximate Connectivity Matrix (using order 1) | $\left( \begin{matrix} -1 & -0.99 & 0 & 0 & 0 & 0.12 \\ -0.71 & -1 & 0 & 0 & 0 & -0.16 \\ 0 & -0.27 & -1 & -0.85 & 0 & 0.16 \\ 0.02 & 0.61 & -0.89 & -1 & 0 & -0.35 \\ 0 & 0 & 0 & -0.71 & -1 & -0.66 \\ 0.01 & 0.01 & 0 & 0.27 & -0.28 & -1 \end{matrix} \right)$  c : Approximate Connectivity Matrix (using order 2) |

Exact connectivity matrix : file "Solution_6K.rda".

d ($r^{e},r)$ = 0.8711854	Order 1

d ($r^{e},r)$ = 0.03618133	Order 2

#### 2.5 Drawing of these networks

Network drawings are based on data corresponding to the order 1 approximate connectivity matrix, utilizing two replicas. In Figure 5 of the article, edges with connectivity coefficients less than 0.1 in absolute value (default parameter of the "DrawGraph" program) are excluded.

| $\left( \begin{matrix} -1 & 0.001 & -0.311 \\ 2.873 & -1 & 0.106 \\ 0.020 & 1.065 & -1 \end{matrix} \right)$  a : "3 kinases" network | $\left( \begin{matrix} -1 & -0.498 & -0.296 \\ -0.208 & -1 & -0.111 \\ -0.400 & -0.474 & -1 \end{matrix} \right)$  b : "Linear 3 nodes" network |
| --- | --- |
| $\left( \begin{matrix} -1 & -0.769 & 0.194 & 0.398 \\ 0.017 & -1 & 0.177 & 0.330 \\ -0.273 & -0.011 & -1 & 0.076 \\ 0.206 & 0.143 & 1.289 & -1 \end{matrix} \right)$  c : "4 nodes" network | $\left( \begin{matrix} -1 & -0.966 & 0.010 & 0.002 & -0.024 & 0.097 \\ -0.646 & -1 & 0.017 & 0.029 & -0.020 & -0.167 \\ 0.046 & -0.101 & -1 & -0.928 & 0.025 & 0.304 \\ -0.202 & 0.203 & -0.846 & -1 & 0.011 & -0.245 \\ 0.045 & 0.055 & 0.098 & -0.739 & -1 & -0.279 \\ 0.217 & 0.433 & 0.199 & 0.522 & -0.347 & -1 \end{matrix} \right)$  d : "Cascade MAPK" (6 noeuds) |

### 3 Simulated networks

#### 3.1 Dream Challenge 4 networks

##### 3.1.1 Getting the data

<https://www.synapse.org/#!Synapse:syn3049712/wiki/74630> .

Data are available for download (files "DREAM4_InSilico_Size10.zip" and "…_Size100.zip").

In each zipped package, there is a folder corresponding to each of the five networks ("InSilico_size10_1" etc.). In these folders, files called "…. _wildtype.tsv" correspond to measurements of the non-perturbed networks, files "… _knockouts.tsv" and "… _knockdowns.tsv" to measurements where a -100% ("KO") or -50% ("KD") perturbation is applied successively to different network nodes.

Solutions are no longer available on the Dream Challenge site now, but the corresponding files ("DREAM4_GoldStandard_InSilico_Size10_1.tsv" etc.) have been archived on our "github" site.

For ease of reference, both data and solutions (formatted for "MRARegress") are accessible on the site <https://github.com/J-P-BORG/MRARegress>, folder "data" (MatExp_10_1.rda to MatExp_10_5.rda and Solution_10_1.rda to Solution_10_5.rda, ditto for 100 node networks).

##### 3.1.2 Standard deviation of "Distances to Diagonal" of the networks

**Supplementary table 1 :** Standard deviation of "Distances to Diagonal" of DC4 networks, depending on the percentage of known data (10 simulations each time)

| % known values | 20 | 40 | 60 | 80 |
| --- | --- | --- | --- | --- |
| InSilico_10_1 | 0.101 | 0.064 | 0.053 | 0.049 |
| InSilico_10_2 | 0.073 | 0.065 | 0.092 | 0.067 |
| InSilico_10_3 | 0.082 | 0.089 | 0.135 | 0.054 |
| InSilico_10_4 | 0.106 | 0.083 | 0.083 | 0.052 |
| InSilico_10_5 | 0.090 | 0.115 | 0.119 | 0.085 |
| InSilico_100_1 | 0.050 | 0.040 | 0.026 | 0.026 |
| InSilico_100_2 | 0.032 | 0.030 | 0.018 | 0.022 |
| InSilico_100_3 | 0.051 | 0.042 | 0.034 | 0.019 |
| InSilico_100_4 | 0.030 | 0.025 | 0.017 | 0.023 |
| InSilico_100_5 | 0.030 | 0.044 | 0.040 | 0.244 |

#### 3.2 FRANK generated networks

##### 3.2.1 Generation of FRANK networks

Numerous networks were generated by FRANK generator (Carré *et al.*, 2017). To launch it: <https://m2sb.org/?page=FRANK>. While the corresponding server may be inactive, contacting the authors of the quoted article can facilitate its reactivation.

This generator delivers a file (.csv) named ""Frankxxxxnonmodified_network.csv" (where xxxx is a number, delivered by FRANK, identifying the file), according to different parameters which are described in § 3.2.2.

This generator simulates genes behavior: genes are regulated. TF act on other TFs and TAs, but TAs don't act on TFs.

In our program, nbN (number of nodes) = TF+TA (TF represents the "number of Transcription Factors" and TA the "number of targets": their outdegrees are 0).

If TA=0, the file matches with a square matrix [nbN, nbN]. If TA>0, we get a rectangular matrix (nbN rows, TF columns). In this case, we add TA null columns on the right, to get a square matrix [nbN, nbN], which is mandatory to use MRA.

Then, we replace the diagonal elements with -1, to comply with MRA requirements. This file represents the "Solution" used to score the results (called ${"r}^{e} "$ in the article).

Starting from the "Solution", we compute the "Global Response Matrix R", where

$$R[i,j]= 2*\frac{X_{i}\left( P0+\Delta Pj \right)-X_{i}\left( P0 \right)}{X_{i}\left( P0+\Delta Pj \right)+X_{i}\left( P0 \right)}$$

We can demonstrate that $R= {{(r}^{e})}^{-1}\left( \text{diag}\left( {{(r}^{e})}^{-1} \right) \right)^{-1}\text{diag}(R)$.

$\text{diag}(R)$ is known since we applied known perturbations to $X_{i}$ concentrations. For simulating full KOs we apply -100% to each $X_{i}$ and so, all the diagonal elements of $\boldsymbol{R}$ are equal to -2, because $X_{i}\left( P0+\Delta Pi \right)=0$. For a KD at -50% they are all equal to -2/3. Refer to (Borg *et al.*, 2023).

Additive $N(0,\sigma)$ noise was added to $\boldsymbol{R}$ with $\sigma=k\bar{X}$, $k$ a factor adjusting the noise level (k=0.1 : "medium noise" and k=0.5 : "loud noise") and $\bar{X}$ the average concentration of the genes.

##### 3.2.2 Simulated FRANK networks

We generated five independent networks for each set, utilizing different sets of TF and TA values.

Simulated measurements files, denoted "Frank_TFxx_TAyy_z_ R1.rda" (medium noise), "Frank_TFxx_TAyy_z_ R2.rda" (loud noise) and "Frank_TFxx_TAyy_z_ Sol.rda" (solution), where (xx.yy)$\in${(30.0),(60.0),(100.0), (30.30), (50.50), (100.100)} and $z\in\left⟦ 1,5 \right⟧$, are located in the "data" folder on our site <https://github.com/J-P-BORG/MRARegress>.

The parameters used are as follows:

- Number of Transcription Factors	: TF number (xxx in the file name)
- Number of TArget genes	: TA number (yyy in the file name)
- Number of eigenvalues of the TF matrix on the unit circle	1	(to get a system leading to a steady state, so as to comply with MRA requirements)
- Value of the variance of the noise added to the log-normal observations : 0 (default value, meaning no noise added by FRANK)
- Time-serie observations	: not concerned. Choose dynamic
- nb expce	: not concerned. Choose 1
- Seed of the random value generator	: define a different value for each of the 5 files. Seeds = c(12345, 72090, 87577, 45648, 16637). These seeds are also used as "seed" for the noise generator.
- Minimum sparsity (number of nonzero elements per row) of the TF and TA matrices	: set to 0.15*(TF+TA), to have a sparsity of 15%
- Maximum sparsity (number of nonzero elements per row) of the TF and TA matrices	: set to 0.15*(TF+TA)+1, to have a sparsity of 15%
- Slop for the sparsity of the matrices	: -2 (default value)
- Magnitude of deviation from zero of the nonnull elements of the matrix	: 1	(default value)
- Value of the mean of the log-normal probability distribution of the observations : 5 (default value)
- Value of the variance of the log-normal probabilty distribution of the observations : 3.5 (default value)

### 4 MRARegress

From gene expression data measurements post-perturbations, "MRARegress" (version 1.0.0) provides all presented results, including the connectivity matrix ("r"), ANOVA data, second-order Taylor coefficients, whether the perturbations are independent or not, if the system rank is sufficient (which it checks), with replicas or not, possibly leading to a system of over-determined equations.

#### 4.1 Precise version of R, R Studio, packages and hardware used

R version 4.3.2 (2023-10-31 ucrt) -- "Eye Holes" - Platform: x86_64-w64-mingw32/x64 (64-bit)

RStudio 2023.06.0 Build 421.

**Modules required by "MRARegress":**

| **Package** | **Version** |
| --- | --- |
| BiocManager | 1.30.22 |
| CVXR | 1.0-12 |
| devtools | 2.4.5 |
| dplyr | 1.1.4 |
| glmnet | 4.1-8 |
| magrittr | 2.0.3 |
| minet | 3.58.0 |
| randomForest | 4.7-1.1 |
| RCy3 | 2.20.2 |
| rootSolve | 1.8.2.4 |
| stats | 4.3.2 |
| stringr | 1.5.1 |
| testthat | 3.2.1 |
| upstartr | 0.1.2 |
| utils | 4.3.2 |

"minet" requires the package "Biocmanager". Then, BiocManager::install("minet").

The software "Cytoscape" (version >= 3.6.1 required -- our version is 3.10.2) must be installed and running before using "DrawGraph", a module of "MRARegress" used to draw network graphs.

**Modules required by "AdvancedMRA.R" (the program used to generate the results of this article):**

| **Package** | **Version** |
| --- | --- |
| CVXR | 1.0-12 |
| data.table | 1.15.2 |
| MRARegress | 1.0.0 |
| RCy3 | 2.20.2 |
| stringr | 1.5.1 |
| upstartr | 0.1.2 |

**Our Hardware:**

PC Dell, core i7 - 14 cores, 2.4 GHz - RAM 64 GB.

**Our code:**

- "MRARegress", available from GitHub: <https://github.com/J-P-BORG/MRARegress>
- "AdvancedMRA.R", available from GitHub: <https://github.com/J-P-BORG/MRA>
- Data files, available from GitHub: <https://github.com/J-P-BORG/MRARegress>

then, folder "data".

#### 4.2 Installation and Basic Testing

- Check the installed versions of R and RStudio. Update them if necessary.
- Check that the required modules (see previous paragraph) are installed and that the versions match, otherwise install and/or update them.
- Install the "MRARegress" package: devtools::install_github("J-P-Borg/MRARegress").
- Check that there is no error message. There may be warnings about replacing the previous import of certain objects.
- The "MRARegress" package should now appear in RStudio's package list (at the bottom right of the screen).

**Two simple examples, to test installation** :

library("MRARegress")

library("testthat")

1^st^ example :

MatExp <- matrix(c(1:12), nrow=3)

MRARegress(MatExp)$r

The following results matrix should be displayed:

```
           Q1        Q2         Q3
N1 -1.0000000  2.954545 -2.0454545
N2  0.3021978 -1.000000  0.7417582
N3 -0.3703704  1.296296 -1.0000000
```

2^nd^ example :

Run "Cytoscape"

```
MatExp	<- matrix(c(1,2,3, 11,21,31, 12,22,32, 13,23,33),nrow=3)
Ret	<- MRARegress (MatExp, Relative=FALSE)
RetDG	<- DrawGraph (Ret)
test_that("Simple network", {
     expect_equal (RetDG$Variables[[2]], 3, tolerance=1E-4)			# 3 nodes (DrawGraph succeeded)
})
```

The following line should be displayed:

Test passed

When entering "Ret$r", the following connectivity matrix should appear:

```
> Ret$r
     Q1 Q2   Q3
N1 -1.0  2 -1.0
N2  0.5 -1  0.5
N3 -1.0  2 -1.0
```

When opening "Cytoscape", one must see:

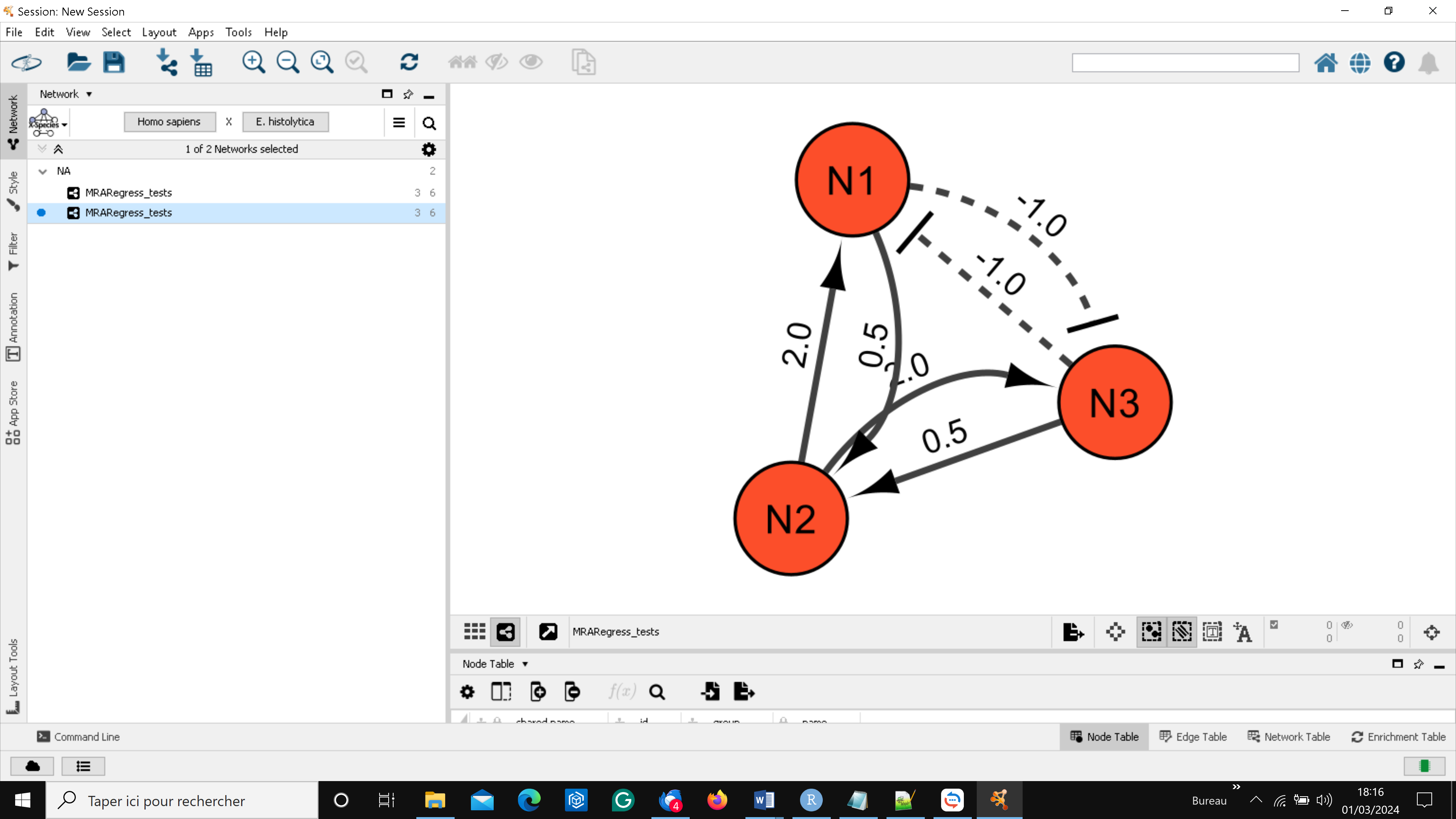

Connectivity matrix values are graphically represented.

In Cytoscape, users can manipulate nodes and arcs to enhance diagram clarity: click on a node to select it, then move it with the mouse. Select an arc by holding down the "alt" key. A red square is drawn next to this arc. Move this square to distort the arc. See "Cytoscape" documentation.

#### 4.3 Results displayed

Depending on user preference, methods such as "Mutual Information" (ARACNE, CLR, MRNET), "Linear Regression" (simple: LSE^[1]^, TLR^[2]^, with restrictions: LASSO^[3]^, or subset selection: STEP^[4]^), "Polynomial Regression "(Order 2), or "Machine Learning" (Random Forest) are applied to the data, considering knowledge contributions if TLR is used.

Various outputs are available (see the package's "thumbnail", available on github), including, for example, the "Connectivity Matrix" 'r', ANOVA results, 95% Confidence Interval of coefficients $r_{i,j},$ a "heat map" of these coefficients, a network description in "GraphML" format, and automatic network drawing valued by connectivity coefficients using Cytoscape. In this depiction, nodes with unsatisfactory linear models are highlighted.

Two methods of connectivity coefficient classification are proposed (threshold-based or quantile-based). Multiple classes, including digitization {0, 1}, can be defined. For known exact connectivity coefficients (" $r^{e}$ "), several comparison scores are suggested to assess " $r$ " and " $r^{e}$ " matrices, including 'Confusion Matrix', 'Precision', 'Recall', 'F1 score', 'Sensitivity', 'Specificity', and 'Distance to Diagonal'. Additionally, a network drawing displaying discovered edges, false positives, and false negatives is available.

Upon loading, users can access user manuals for each module comprising the "MRARegress package":

MRARegress:	the main module that performs all the calculations mentioned in this	article (connectivity matrix, ANOVA …).

rCI:	computation of $r_{i,j}$confidence intervals.

DrawHeat:	"heat map" drawing.

DrawGraph:	valued network drawing.

DrawGraphM:	network description in 'GraphML' format.

Classify:	discretization of connectivity coefficients.

Score:	computation of various scores.

DrawDiscr :	network drawing, with actual edges discovered, false positives and false	negatives.

### 5 Références

Alon,U. (2006) An Introduction to Systems Biology: Design Principles of Biological Circuits Chapman and Hall/CRC, New York.

Borg,J.-P. *et al.* (2023) Modular response analysis reformulated as a multilinear regression problem. *Bioinformatics*, **39**, btad166.

Carré,C. *et al.* (2017) Reverse engineering highlights potential principles of large gene regulatory network design and learning. *NPJ Syst. Biol. Appl.*, **3**, 17.

Sontag,E. *et al.* (2004) Inferring dynamic architecture of cellular networks using time series of gene expression, protein and metabolite data. *Bioinforma. Oxf. Engl.*, **20**, 1877–86.

Thomaseth,C. *et al.* (2018) Impact of measurement noise, experimental design, and estimation methods on Modular Response Analysis based network reconstruction. *Sci. Rep.*, **8**, 16217.

1. LSE : Least Square Estimation (choose "TLR method" to use this cost function).
2. TLR : Threshold Linear Regress. This method we developed (§ 1) reaches very good results, depending on noise and network size. Processing time is much lower than that of LASSO or STEP methods (Borg *et al.*, 2023).
3. LASSO : Least Absolute Shrinkage and Selection Operator
4. STEP : Forward and Backward Stepwise Selection
